# Microbial Invasion and Immunosuppression Drives Adenoma Progression in Early Colorectal Cancer Development

**DOI:** 10.64898/2026.07.10.737736

**Authors:** Wei Liang, Lisa Falk, Daniele Lucarelli, Philipp Putze, Amira Metwaly, Yiran Zheng, Fabian Springer, Thomas Winogrodzki, Qixia Chan, Theresa M. Pauli, Yue Zhang, Georg Zeller, Matthias Meier, Angelika Schnieke, Friederike Ebner, Dirk Haller, Tatiana Flisikowska, Dieter Saur, Krzysztof Flisikowski

## Abstract

Several inherited predispositions to colorectal cancer exist, most notably familial adenomatous polyposis (FAP), which typically involves a germline loss-of-function mutation in one *APC* allele. A second hit in *APC* or related loci leads to aberrant Wnt signalling, triggering adenoma formation with near-complete penetrance. Yet, substantial variability in disease onset and severity, even among siblings with identical *APC* germline mutations, implicates environmental modifiers. Emerging evidence points to the gut microbiome as a critical regulator of adenoma initiation and progression, particularly in early-onset CRC. Here, we show that bacterial invasion is associated with neutrophil immunosuppression, T-cell exclusion and adenoma progression in a porcine model of FAP. Longitudinal mapping of progressing and regressing polyps using single-cell RNA sequencing, spatial transcriptomics and integrated microbiome profiling resolved the cellular and microbial architecture underlying these divergent lesion states. Single-cell RNA-seq identified 35 cell subpopulations and showed that regressing polyps are enriched for central-memory CD4+ T cells, cytotoxic CD8+ T cells and inflammatory neutrophils, whereas progressing polyps contain immunosuppressive/PD-L1-associated neutrophils, regulatory and dysfunctional T cells. Spatial transcriptomics with a custom host panel and targeted bacterial probes linked bacterial-high adenomatous regions to immunosuppressive neutrophil niches and T cell exclusion, while regressing lesions showed predominantly lumen-adjacent bacterial localization and spatially organized immune surveillance. These results provide a spatially resolved map of cellular and microbial organisation in early colorectal polyp progression and regression and reveal a microbiome-driven, neutrophil-mediated mechanism of early tumour progression with implications for CRC interception.

## Introduction

Colorectal cancer (CRC), progresses in a stepwise fashion from a benign polyp to metastatic disease through a series of well-defined genetic and epigenetic alterations, a process that may take decades ^1^. This long CRC timeline creates a substantial window of opportunity for preventative intervention from suppressing tumour initiation (primary prevention), preventing tumour progression (secondary prevention), or management of chronic disease (tertiary prevention) ^2^. For decades, the concept of immune surveillance, postulating that a normal cell acquiring oncogenic mutations can be recognised as foreign and eliminated by the immune system ^3^ has been appreciated as a critical component of CRC prevention ^4^. This theory is supported by the observation that CRC progression is associated with changes in the composition of tumour infiltrating cells, which create an immuno-suppressive microenviroment^5^.

Polyps are considered precursor lesions to CRC. For a polyp to become malignant, it must accumulate multiple genetic mutations and epigenetic changes, a process that takes several years. This extended timeline allows for preventive colonoscopy screening and treatment, which significantly reduces CRC mortality. During colonoscopy, polyps are classified as either low-(smaller than 10 mm) or high-risk adenomas, primary based on size. The need for surveillance for small adenomas remains a subject of debate. Prospective study showed that around 25% of small adenomatous polyps (< 9 mm) progressed to invasive CRC within 5 years ^6^, the fate of the remaining polyps is less clear. Several studies have reported a spontaneous regression of small polyps in humans ^7–11^.

The term spontaneous regression is obviously misleading, as there is likely a causative factor. Gaining insights into the natural mechanisms behind polyp regression may open the way for more effective therapies and prophylactics. Due to the longitudinal nature of this process, studying it in human patients is difficult.

We generated pigs carrying the *APC^1311/+^* mutation, which is orthologous to the human *APC^1309^* hotspot mutation associated with severe FAP in humans ^12^. In our previous studies, we showed that these pigs recapitulate major hallmarks of the human disease ^13^, show variable polyposis severity ^14^, and provide a translational model for preclinical studies ^15–17^. Moreover, disease progression was influenced by a Western diet ^17^.

In this longitudinal study, we characterise the dynamic trajectories of colon polyps in a large cohort of *APC^1311/+^* pigs, capturing both progressive and regressive polyp states. We further demonstrate that polyp regression is orchestrated by spatially coordinated interactions between the gut microbiome and the local immune system, with microbial niches guiding neutrophil activation and cytotoxic lymphocyte engagement to drive lesion clearance.

## Material and Methods

### Ethics statement

Animal experiments were approved by the Committee on Animal Health and Care of the local government body of the state of Upper Bavaria (Permission No. 55.2-2532.Vet_02-18-33) and performed according to the German Animal Welfare Act and European Union Normative for Care and Use of Experimental Animals.

### Pig housing

The *APC^1311/+^* pigs (German Landrace x Minipig) were housed at the TUM Animal Research Center (ARC). Piglets were weaned at five weeks. All animals received the same diet (HEMO U134 pellets (Likra West)) and water ad libitum.

### Tissue sampling

Normal mucosa (NM, n= 30) and polyp (n= 60) samples were collected from *APC^1311/+^* pigs (n= 30) and stored at −80°C for further molecular analyses. For histological and immunohistochemical analyses tissue sections were rinsed in cold phosphate-buffered saline (PBS) and fixed in 4% paraformaldehyde (Sigma-Aldrich, USA) for 24 hours at room temperature. After fixation, tissue specimens were transferred into 70% ethanol and then embedded in paraffin.

### DNA extraction

DNA isolation from tissues and cells was done with QuickExtract™ DNA Extraction Solution (Biosearch Technologies).

### RNA isolation

RNA was isolated from bulk tumour samples and cancer cell cultures using a Monarch Total RNA Miniprep Kit (NEB) according to the manufacturer’s protocols.

### RT-PCR

A total of 400 ng of RNA was used for the cDNA synthesis using LunaScript RT Master Mix Kit (NEB) according to the manufacturer’s protocol. Diluted cDNA (1:3) was used for RT-PCR (GoTaq® G2, Promega). The resulting products were visualised after 2% agarose gel electrophoresis.

### QRT-PCR

Real-time quantitative PCR (qPCR) was performed using the Qpcrbio SyGreen Mix Lo-ROX (Biosystems) in a QuantStudio 5 Real-Time-PCR-Cycler (Thermofisher Scientific) with default thermal cycling parameters. Reactions were performed in 10 μl volume. Samples were assayed in triplicate and relative expression from the bulk tumour samples was normalised to *EIF2B1* expression. Relative expression was calculated using the ΔΔCt method. One-way ANOVA was performed, followed by Tukey’s test for statistical evaluation using GraphPad Prism 8. The list of primers used for qPCR is shown in **Table S1**.

### Digital droplet PCR (ddPCR)

The *APC* copy number was determined by ddPCR. Genomic DNA was digested with 3 U /μg of HindIII (NEB, Frankfurt am Main, Germany). TaqMan PCR reaction (23 μL final volume) was set up using 100 ng digested DNA, 2x ddPCR supermix for probes (BioRad Laboratories, Herkules, USA), 20x target primer/FAM-labeled probe, and 20x reference primer/HEX-labelled primer probe (Eurofins, Germany). Droplets were generated using a QX200 Droplet Generator combining 20 μL TaqMan PCR with 70 μL droplet generator oil for probes in a DG8 Cartridge and then transferred to PCT Plate 96 (Eppendorf, Germany). PCR was performed with the following cycling conditions: 95°C for 10 min., followed by 40 cycles of 94°C for 30 sec., 61°C for 1 min., and a final hold at 98°C for 10 min. The proportion of PCR-positive to PCR-negative droplets was determined using a QX200 droplet reader, and data were analysed using QuantaSoft Software (BioRad Laboratories, Herkules, USA).

### Pyrosequencing

Pyrosequencing was used to measure the DNA methylation in the promoter regions of target genes. Pyrosequencing assays were designed using PyroMark Assay Design 2.0 software (Qiagen).

500 ng genomic DNA was bisulphite-converted with the EZ DNA Methylation-Direct kit (Zymo Research, Irvine, USA) according to the manufacturer’s instructions. PCR samples were amplified using PyroMark PCR kit (Qiagen). PCR products were sequenced using PyroMark Q48 Advanced CpG reagents on a PyroMark Q48 Autoprep instrument (Qiagen).

### Immunostaining

Tissues samples were fixed with 4 % paraformaldehyde for 24 h and store at 70 % ethanol. The samples were dehydrated and embedded in desired orientation with paraffin wax. Sections from them were cut at a tickness of 4 μm and deparaffinised before antigen retrieval. For frozen sections, tissues were embedded in OCT and stored at −80 °C. Sections were prepared at −20 °C and air dried before fixation and staining.

Staining was carried out with the prepared sections. Sections were blocked with 0,5% serum (from the animal of secondary antibody) in PBS followed by 1 h incubation of corresponding primary antibody and 1 h incubation of secondary antibody.

The Ki67 expression was examined by immunohistochemistry using Ki-67 antibody (DSC innovative Diagnostik-System KI681C002, dilution 1:300) with secondary antibody Goat Anti-Rabbit IgG-HRP (Santa Cruz sc-2780, dilution 1:400). The CD3+ cells were detected by CD3 antibody (Mouse, Southern Biotech 4511-01, dilution 1:100) with secondary antibody m-IgGκ BP-Biotin (Santa Cruz Biotechnology, sc-516142, dilution 1:200), VECTASTAIN® Elite® ABC Kit (Vectorlab PK-6100) was used to bring in peroxidase (HRP) for color developing. MPO and IBA1 were detected using anti-myeloperoxidase antibody (Dianova DLN-012930, dilution 1:300) and an anti-IBA1 (Wako FujiFilm 019-19741, dilution 1:1000) antibody, respectively. Sections were subsequently incubated with a goat anti-rabbit IgG-HRP the secondary antibody (Vector Laboratories BA-1000, dilution 1:200). HRP activity was visualised using diaminobenzidine (DAB) as the chromogenic substrate.

Positive staining was quantified by analysing ten randomly selected fields (magnification x40) from each tissue section. Data are presented as the percentage of positevely stained cells among all cells counted in each section. Statistical significance was calculated using t-test. * P< 0.05, ** P< 0.01, *** P< 0.001.

### Laser microdissection

Laser microdissection (LMD) of cryosectioned samples was performed immediately after H&E staining using a Leica Microsystems Laser Microdissection Systems 6000 and Leica Application Suite software (Leica). In total, ten LMD-captured crypts per normal mucosa sample were cut and collected into 50 µL lysis buffer from the AllPrep® DNA/RNA Micro Kit (Qiagen). Dissected samples were stored at −80°C. Total RNA was isolated using the AllPrep® DNA/RNA Micro Kit according to the manufacturer’s instructions (Qiagen).

### Isolation and cultivation of porcine intestinal crypts

Epithelial crypts were isolated from regressing and progressing polyps from *APC^1311/+^* pigs. Luminal contents were removed from colon by scraping with forceps and washing in 80% EtOH and PBS. The polyps were incubated for 20-25 min under constant shaking in dissociation buffer containing 30 mM EDTA and 10 mM DTT. Then, the samples were washed in PBS and transferred to Advanced DMEM/F12 medium (Corning, USA) supplemented with 10 µM thiazovivin and 100 µg/mL primocin. Crypts were mechanically dissociated with syringe needles under a stereo microscope. We embedded 350-500 crypts per 50 uL of Matrigel in a 24-well plate and incubated for 15 minutes at 37°C, 5% CO_2_. After solidification, the culture was overlaid with 500 uL of growth medium supplemented with 50% murine Wnt3a conditioned medium (equivalent to 50-60 ng/mL recombinant), 4% human R-spondin conditioned medium (equivalent to 2.4 µg/mL recombinant human protein), 1.3% porcine Noggin conditioned medium (equivalent to 100 ng/mL recombinant human Noggin), B27 and N2 supplements, 10 mM nicotinamide, 50 ng/mL recombinant human EGF, 10 nM L-Gastrin, 500 nM LY2157299, 10 mM SB202190 and 2.5 µM CHIR99021, 10 µM thiazovivin and 100 µg/mL primocin. After two days of culture, thiazovivin was removed from the medium.

### Organoid cultivation

The organoids were incubated at 37°C, 5% CO_2_ in a freshly prepared growth medium described above. Two days before passaging, the growth medium was supplemented with 10 µM thiazovivin. Organoids were harvested by removing the medium and resuspending the Matrigel drop in 1 mL of ice-cold PBS. Matrigel was removed by centrifugation at 300 x g for 5 minutes at 4°C and the cells were dissociated for 10 minutes at 37°C in TrypLE Express (Thermo Fisher, Cat.# 12604013) and mechanically by pipetting up and down for 10-20 times through a BSA-precoated blunt-end needle. The treatment was stopped by adding the same volume of Advanced DMEM/F12 medium supplemented with 5% foetal calf serum (FCS). The concentration of single cells was determined with a Neubauer chamber. Organoids were passaged weekly and 1-5 x 10^4^ cells were aliquoted per 50 µL of Matrigel in a 24-well plate in six technical replicates. 100 organoids from progressing and regressing polyps were cultivated in parallel and their morphology was assessed after 2 days in culture. Organoids were counted on bright field images taken for each of the technical replicates (n= 6). Images were adjusted for contrast and brightness and loaded into Cellpose^18^. The mask created by the segmentation algorithm was loaded in Fiji/ImageJ and processed by the MorphoLibJ Plugin for Morphological Segmentation. The watershed lines were subjected to the option ‘Analyse Particles’, which counted single particles with a minimum particle area of 300 µm^2^. Data were collected and processed in RStudio.

### Organoids viability staining

Organoids were stained in Matrigel. Staining solution contained 8 ug/mL fluorescein diacetate and 20 ug/mL propidium iodide in PBS. Medium was removed and the wells were washed once with 1 x PBS. The Matrigel drop was covered with staining solution (approximately 500 uL) and incubated at RT for 5 min in the dark. Staining solution was removed and the well was washed twice with 1x PBS. The culture was overlaid with 1x PBS and fluorescent images were taken.

### 16S rRNA Gene Amplicon Sequencing and Microbiome Analysis

Total genomic DNA was extracted from faecal samples of *APC^1311/+^* pigs aged 5.5 ± 0.8 months using a bead-beating protocol, followed by 16S rRNA gene amplicon sequencing as previously described^19^. Bioinformatic processing was performed in R v4.2.1 using the Rhea pipeline^20^, and zOTU taxonomic identity was verified against the EzBioCloud 16S database (v2023.08.23)^21^. Raw zOTU counts were normalised by proportional min-sum scaling to the minimum library size across all samples. Alpha diversity was characterised by the Shannon effective number of species (exp H′), computed on the full normalised zOTU table; between-group differences across normal mucosa (NM), Progressing, and Regressing tissue types were tested by the Kruskal–Wallis rank-sum test. Beta diversity was assessed using Bray– Curtis dissimilarity visualised by non-metric multidimensional scaling (NMDS); group separation was tested by PERMANOVA (adonis2, 999 permutations). Phylum-level relative abundances were aggregated from genus-level profiles and expressed as mean per group. Differential abundance between tissue types was tested by the Wilcoxon rank-sum test with Benjamini–Hochberg false discovery rate correction; only taxa reaching p < 0.05 are shown, ranked by absolute difference in mean relative abundance (%), and effect sizes are additionally expressed as log₂ fold change.

### Pig polyp Sample Preparation for sc-RNAseq

Fresh polyp samples were minced and enzymatically digested with the human tumour dissociation kit (Miltenyi #130-095-929) in DMEM medium (Sigma, #D5796-500 mL) for 45 min at 37°C with agitation. The cell suspension was strained through a 100 μm strainer, pelleted and resuspended in 2% FCS/PBS. Cell debris was removed using Debris removal solution (Miltenyi, catalog #130-109-398) and live cells were enriched using the Dead Cell Removal Kit (Miltenyi Biotec, #130-090-101), both according to the manufacturer’s instructions.

### sc-RNAseq Library Preparation and Sequencing

Single-cell GEM generation, barcoding, and library construction were performed using the 10x Genomics Chromium Single Cell 3’ Reagent Kits (v3.1 Chemistry) according to the manufacturer’s instructions. Cells were counted, diluted in 2% FCS/PBS, and up to 30,000 cells were loaded per lane on a Chromium Next GEM Chip G to generate gel beads in emulsion. cDNA and libraries were assessed for quality and fragment size using an Agilent Tapestation 4200 using DNA HS 5000 tape. Libraries were sequenced on the Illumina NovaSeq 6000 S2 (PE; 28+94 bp).

The second experiment was conducted with the Whole Transcriptome Analysis (WTA) Amplification Kit from BD Rhapsody™ according to the manufacturer’s instructions. For each sample, 30,000 cells were captured and sequenced on the Illumina NovaSeq X.

### Alignment with *Sus scrofa* 11.1 Reference genome

FASTQ files obtained from the 10x Genomics Chromium experiment were processed with Cell Ranger v9.0.1 (10x Genomics) using cellranger count with the pig reference (sus_scrofa_GEX_ENS113). The obtained FASTQ sequencing files from the BD Rhapsody experiment were processed via the cloud-based Seven Bridges Platform environment (Seven Bridges Genomics) using the BD Rhapsody Sequence Analysis Pipeline.

The pig reference was built with the *Sus scrofa* 11.1 reference genome and ENSEMBLE 113 annotation file for both experiments. After alignment, ENS identifiers were converted to their corresponding gene symbols where available.

### Data preprocessing and quality control of scRNA-seq Library

Ambient RNA was removed using scAR on the filtered feature barcode matrix using default parameters. Further analysis was done with Python 3.11.14 and Scanpy 1.11.5.

Calculation of quality metrics and subsequent filtering of low quality cells based on median absolute deviations (MAD) was performed in accordance to single-cell best practices. The quality filtering was done in two rounds. Initially the whole dataset was filtered by nMAD of 5, doublets were removed using scrublet with a threshold of 0.1, cells expressing fewer then 200 genes were removed and clusters expressing haemoglobin genes were removed. Samples were integrated using scVI on the intersection of 2.000 highly variable genes across all samples. Raw count matrices from each dataset were normalized using scanpy’s normalize_total function (target sum = 1 × 10⁶) followed by logarithmic transformation (log1p). Dimensionality reduction was performed using Uniform Manifold Approximation and Projection (UMAP) and cells were grouped based on Leiden clustering and marker genes were identified with Scanpy’s rank_genes function using the Wilcoxon test. First-level annotations were assigned using canonical marker genes and comprsed epithelial, stromal, T Cell, B Cell, myeloid and neutrophil comprtments. MAD-based quality filtering was then repeated separately within each first-level compartment.

### Differential Gene Expression and Gene Set Enrichment Analysis (GSEA)

Pseudobulk profiles were generated from single-cell RNA-seq data to enable differential gene expression analysis between experimental conditions. The analysis was performed using the *decoupleR* package. For each sample, cells were randomly shuffled and evenly split into two groups, forming two pseudobulk replicates. The raw counts of each group were summed to generate the pseudobulk profiles. Lowly expressed genes were filtered prior to differential expression testing using the default expression-based filtering strategy. Differential gene expression (DGE) analysis was then performed on the filtered pseudobulk count matrix using the *DESeq2* package. A negative binomial generalized linear model was fitted with experimental condition as the design factor, and Wald tests were used to assess differential expression between conditions. P values were adjusted for multiple testing using the Benjamini–Hochberg procedure. GSEA was conducted using *gseapy* and the GO Biological Processes gene sets.

### Cell state characterisation

To visualize relative shifts in cell state distributions, Scanpy’s *embedding_density* function was used. Gaussian kernel density estimations within the UMAP embedding space were calculated and plotted for each level one compartment, respectively. Scores for individual cell states were calculated with Scanpy’s *score_genes* function. (See table Scores_per_Celltype.xlsx) Scores and gene sets are provided.

### Xenium Spatial Transcriptomics

For each of the 16 samples, a 5 µm section was placed on Xenium slide and Xenium spatial profiling was performed according to the manufacturer’s instructions. The Xenium Cell Segmentation Staining Reagents Kit (10x Genomics, PN-1000661) and a 480-gene custom panel including five custom probes for bacterial detection were used. Fluorescent in situ hybridization (FISH) was performed post-Xenium.

Custom Probes for detecting bacterial signal:

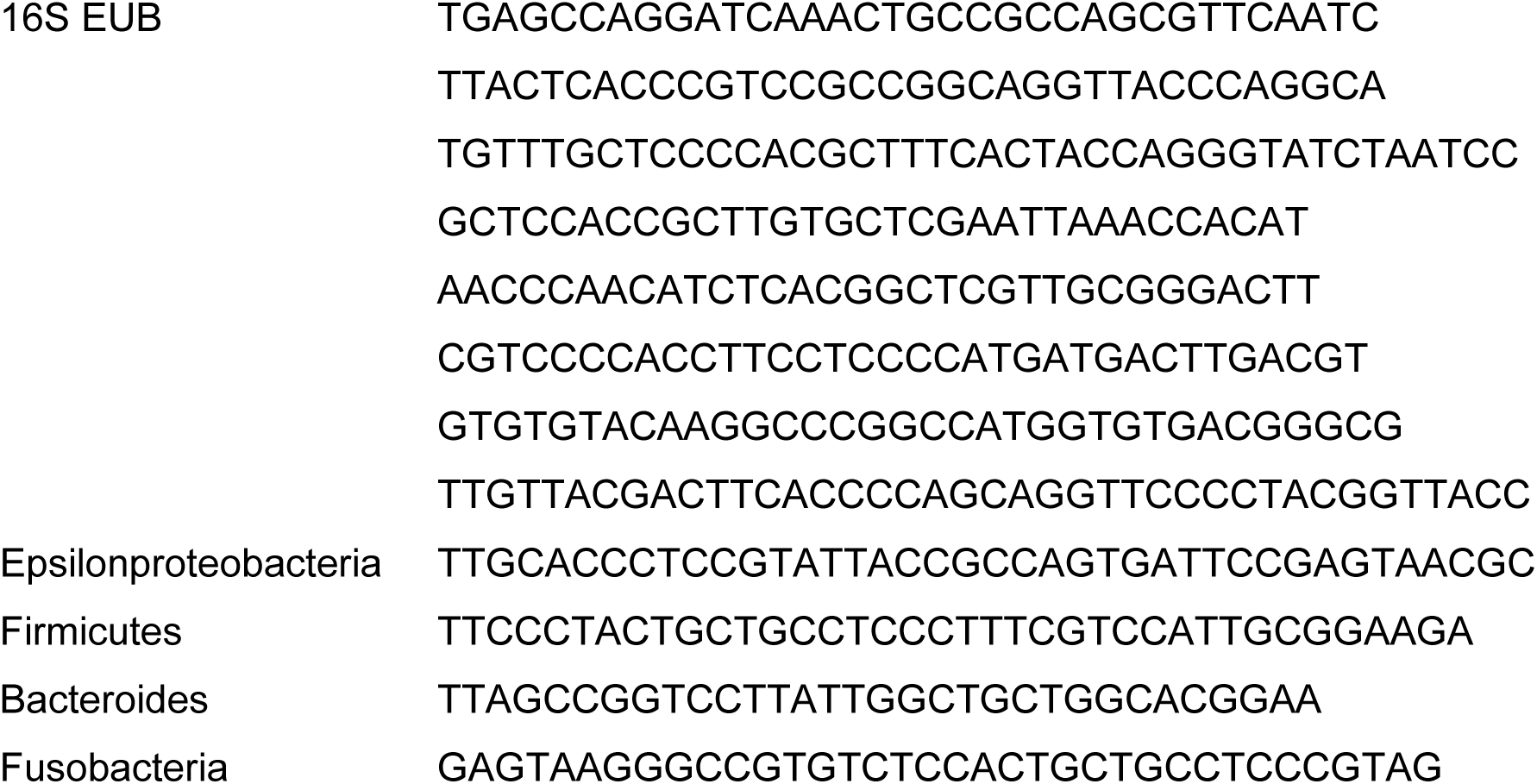

## Results

### Longitudinal characterisation of polyp abundance and phenotype

A total of 82 *APC^13^*^11^*^/+^*mutant pigs spanning three generations underwent longitudinal colonoscopic surveillance at 3, 6, and 9 months of age, followed by extensive multimodal molecular and histopathological analyses (**Figure 1A**). Consistent with our previous observations, ^14^ the initial examination revealed striking inter-individual heterogeneity in tumour burden, with colonic polyp numbers ranging from 9 to more than 300 lesions per animal (**Figure 1B**). To examine whether the severity of polyposis influences subsequent lesion dynamics, animals were stratified according to the median polyp burden into low-polyp (LP; <200 polyps) and high-polyp (HP; >200 polyps) groups. Across all animals, 14,802 polyps measuring 3–6 mm were documented at baseline (**Figure 1B,C**). Despite substantial variability in lesion number, polyps displayed highly uniform endoscopic morphology, and histopathological evaluation of representative lesions (n = 136) identified all as low-grade adenomas.

**Figure 1.**
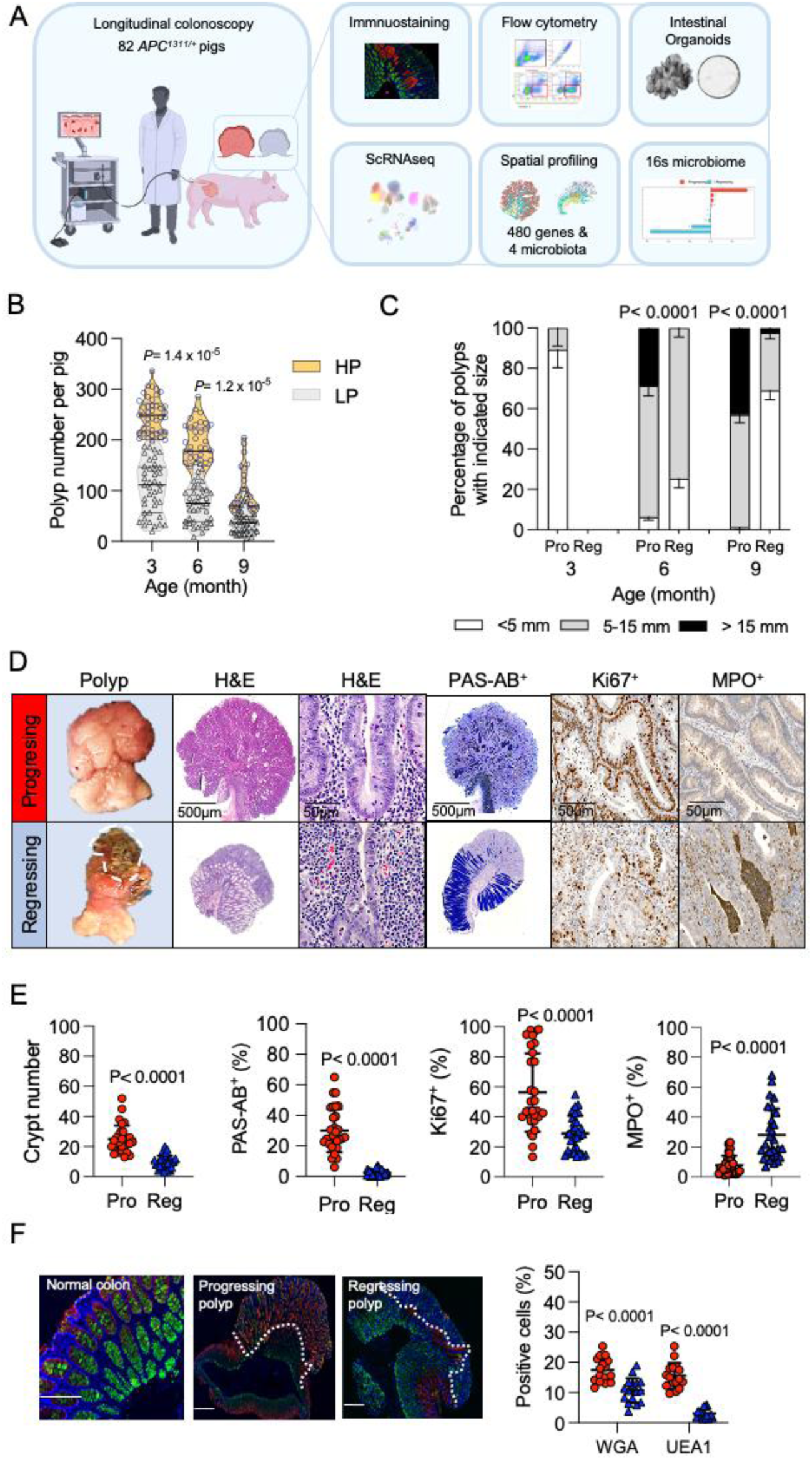
Characterisation of polyp regression. **A.** Overview of study design. **B**. The numbers of colon polyps counted during the colonoscopy the age of 3-, 6-, and 9-months. HP: high polyp pigs (n> 200 at the age of 3-months, n= 54), LP: low polyp pigs (n≤ 200, n= 28). **C**. The percentage of polyps with indicated size. **D**. Representative images showing the morphology, hematoxylin-eosin, and Periodic acid–Schiff and Alcian blue (PAS-AB) stainings, IHC for Ki67^+^ and MPO^+^ in progressing and regressing polyps. **E.** The boxplots show the quantitative measurement for number of crypts, PAS-AB^+^, Ki67^+^ and MPO^+^ in the marginal regions of progressing (n= 30) and regressing (n= 30) polyps. Each dot in the boxplots indicates the mean count from ten microscopic fields assessed at 20x magnification. The data are presented as the mean ± s.d. Statistical significance assessed using Student’s *t-*test. *** *P*< 0.001. **F**. Immunofluorescence of Ulex Europaeus Agglutinin I (UEA1) and Wheat Germ Aglutinin (WGA) for normal mucosa, progressing and regressing polyp. UEA1+, red signal; WGA+; green signal; DAPI, blue signal. Scale bar: 200 μm.

Longitudinal follow-up revealed a progressive decline in overall polyp burden, most pronounced by 9 months of age (**Supplementary Figure 1A**). The total number of assessed lesions decreased to 10,558 at 6 months and to 4,968 at 9 months, with the reduction occurring more rapidly in HP animals than in LP animals (**Supplementary Figure 1B**). Concomitantly, a substantial subset of lesions acquired a distinct morphology characterised by elevated polypoid structures covered by a white surface layer and focal areas of tissue degeneration suggestive of apoptosis (**Figure 1D; Supplementary Video 1**). Histological analysis of these lesions identified three spatially distinct compartments: a basal region containing normal epithelial crypts, an intermediate adenomatous compartment, and a peripheral marginal zone marked by crypt distortion, crypt loss, abscess formation, and extensive immune-cell infiltration (**Figure 1D,E; Supplementary Figure 2A**). On the basis of these features, these lesions were classified as regressing polyps.

To define molecular differences associated with regression, we compared established markers of neoplastic progression between progressing and regressing lesions. Periodic acid– Schiff/Alcian blue (PAS-AB) staining demonstrated an approximately 30-fold reduction in mucin abundance within the marginal regions of regressing polyps relative to progressing lesions (**Figure 1D,E; Supplementary Figure 2B**). This reduction was accompanied by markedly decreased Ki67 expression (**Figure 1D,E; Supplementary Figure 2C**), indicating diminished proliferative activity during regression. In parallel, β-catenin expression was significantly elevated in progressing compared with regressing polyps (49.9% versus 40.8%, P < 0.001), with the largest differences observed within marginal regions, likely reflecting the progressive loss of crypt architecture in regressing lesions (**Supplementary Figure 2D**). Strong β-catenin localisation persisted in adenomatous crypts and in luminal surface epithelial cells. Immunohistochemical analyses further revealed extensive infiltration of MPO+ neutrophils into adenomatous crypts, accompanied by marked accumulation of IBA1+ macrophages (**Figure 1D,E; Supplementary Figure 2E**) and CD3+ T cells (**Supplementary Figure 2F**) within the marginal compartment of regressing lesions.

Lectin-based profiling of mucin composition using WGA (wheat germ agglutinin) and UEA1 (Ulex europaeus agglutinin I) staining further distinguished progressing from regressing lesions (**Figure 1F, Supplementary Figure 3**). In regressing polyps, UEA1+ staining was largely restricted to the basal and intermediate regions, whereas WGA+ signal delineated single-layered epithelial cells and was detected within surface mucus and isolated goblet-cell vacuoles. By contrast, progressing polyps displayed extensive and overlapping UEA1+ and WGA+ staining throughout adenomatous crypts, with prominent accumulation within peripheral mucus and a sharply demarcated UEA1+ boundary separating dysplastic lesions from adjacent normal-like glands.

Caspase-3 staining revealed significantly elevated apoptosis in regressing polyps (11.3% vs. 4.3%, p < 0.05), with prominent accumulation in marginal and regressive crypt regions (**Supplementary Figure 4A)**. This was corroborated by TUNEL and flow cytometry analyses, which showed higher apoptotic indices in regressing lesions, including increased TUNEL-positive cells and a greater proportion of Annexin V–positive cells (∼1.8-fold). Collectively, these findings demonstrate markedly enhanced apoptotic activity and reduced cell viability in regressing polyps compared to progressing lesions (**Supplementary Figure 4B)**.

The frequency of regression differed substantially according to initial tumour burden. HP animals exhibited a significantly greater proportion of regressing polyps than LP animals (**Supplementary Figure 5A**). At 6 months, regressive features were observed in a median of 60% of lesions in HP pigs (range, 3–90%), compared with 45% in LP pigs (range, 0–81%). By 9 months, the proportion of regressing lesions declined modestly in HP animals to 51% (11– 92%), whereas it increased in LP animals to 55% (12–87%) (**Supplementary Figure 5B**). Consequently, by 9 months of age, progressing and regressing lesions were present in approximately equal proportions in HP animals (**Supplementary Figure 5C**). Divergent growth trajectories between lesion states became increasingly evident over time. Although progressing and regressing lesions were comparable in size at earlier stages, by 9 months progressing polyps reached diameters of 8–20 mm, whereas more than 60% of regressing lesions measured less than 5 mm (**Supplementary Figure 5D**). No sex-specific differences in regression frequency were detected (**Supplementary Figure 5E**).

A subset of fifteen animals underwent serial colonoscopic monitoring for up to two years. During this extended observation period, overall polyposis severity remained stable, whereas individual progressing lesions continued to enlarge, reaching diameters of up to 40 mm (**Supplementary Figure 6A, B**).

30 *APC^1311/+^* pigs (summarised in **Supplementary Figure 7)** underwent a detailed postmortem examination of polyposis. Both progressive and regressive polyps were identified throughout the large intestine **(Supplementary Figure 8A)**. The highest numbers of total and regressing polyps were observed in the distal colon and rectum, particularly in the HP group (**Supplementary Figure 8B**). The distribution of polyposis was correlated with mutant *APC* (mut*APC*) mRNA expression, which was 5-fold higher in the distal colon of HP group than in the proximal colon of LP *APC^1311/+^* pigs (**Supplementary Figure 8C**). This finding aligns with Wnt signaling gradient and tumour distribution observed in the human gastrointestinal tract^22^. Taken together, these findings indicate on an intense regression of polyps in the *APC^1311/+^* pigs occurring between 6 and 9 months of age.

### Molecular signatures of polyp regression and progression

Based on the molecular pathways implicated in early CRC tumorigenesis^23^, together with findings showing a positive correlation between mut*APC* mRNA expression and polyp burden (**Figure 2A**), *APC* was identified as a candidate gene associated with the developmental dynamics of polyps. Quantitative PCR, digital droplet PCR and pyrosequencing revealed a modest reduction in mut *APC* mRNA expression (**Figure 2B**), accompanied by an increased *APC* copy number in regressing polyps (**Figure 2C**). In parallel, elevated 3’UTR DNA methylation in the middle and marginal regions of regressing polyps (**Supplementary Figure 9)**. No differences in *APC* expression were detected in whole polyp sample, likely reflecting cellular heterogeneity (**Supplementary Figure 10)**.

**Figure 2.**
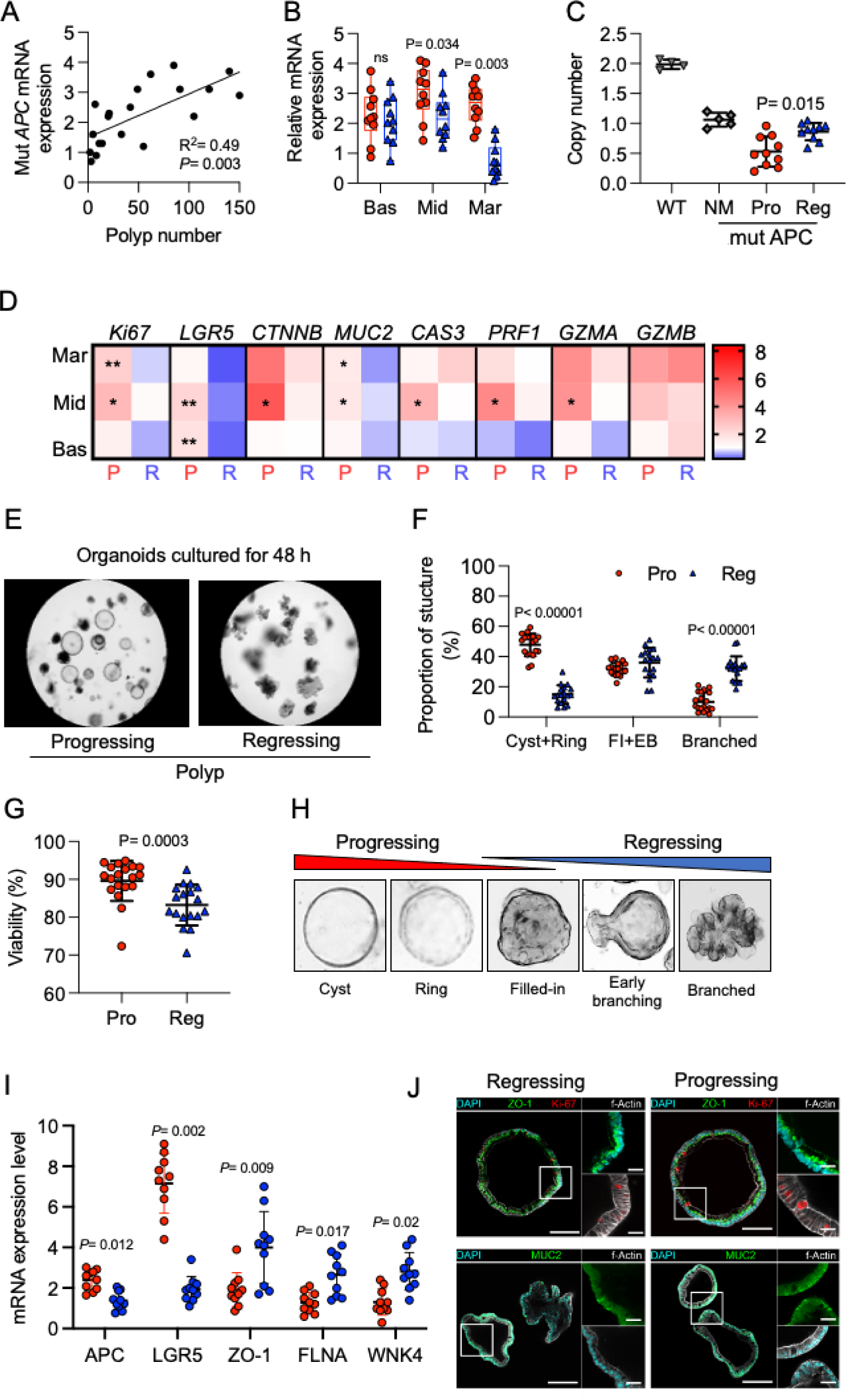
Molecular signatures of regressing polyps. **A.** Plot showing the correlation between the mutant *APC* mRNA expression and number of colon polyps. **B**. mRNA expression of mutant *APC* in different regions of microdissected progressing (n= 10) and regressing (n= 10) polyps. The expression was normalised to normal mucosa. **C**. Copy number of wild-type *APC* in normal mucosa (n= 5), progressing (n= 10) and regressing (n= 10) polyps from *APC^1311/+^* pigs. Wild type pigs (n= 3) were used as a control. ** *P*< 0.01. **D.** Heatmap showing the mRNA expression of selected genes in different regions of microdissected progressing (n= 10) and regressing (n= 10) polyps. Mar, marginal region; Mid, middle region; Bas, basal region. **E**. Representative bright field images of polyp organoids at passage 0 cultured for 48 h. **F**. Quantification of different morphological organoid structures derived from progressing (n= 12) and regressing (n= 12) polyps cultured for 48 h. Note: FI + EB indicate filled-in and early branching organoids. Each dot in the barplot indicates organoid’s structure proportion from one polyp. The data are presented as the mean ± s.d. Statistical significance assessed using two-way anova and multiple comparison. *** *P*< 0.001. **G.** Viability of organoids observed after 48 h of culture. Each dot in the boxplot indicates total number of cystic to branched organoids divided by total number of organoids including burned out. **H.** Overview of organoid structures representing healthy- and disease-like polyp properties. **I.** Expression of genes related to cytoskeleton and colon crypt function (*APC*, *LGR5*) organoids derived from progressive (n= 10) and regressive (n= 10) polyps. **J.** Immunostaining showing the KI67, ZO-1 and MUC2 expression in organoids derived from progressing and regressing polyps. The data are presented as the mean ± s.d. Statistical significance assessed using Student’s *t-*test. ** *P*< 0.01, *** *P*< 0.001.

Collectively, these results indicate that loss of the second *APC* allele represents an important molecular event distinguishing the progressing versus regressing potential of polyps.

In addition, quantitative PCR analysis revealed a differential expression of genes associated with proliferation *(Ki67*), intestinal stem cells *(LGR5),* Wnt signaling *(CTNNB),* the mucus barrier *(MUC2),* and apoptosis *(CAS3, PRF1, GZMA, GZMB)* in different regions of progressing and regressing polyps (**Figure 2D**).

Next, organoids were generated from progressing and regressing polyps to compare their intrinsic epithelial properties (**Figure 2E**). Compared to organoids derived from progressing polyps, rthose derived from egressing polyps showed a significantly higher proportion of branched structures (38% ± 12% vs. 11% ± 9%; *P* < 0.001) and a lower proportion of cystic structures (18% ± 14% vs. 46% ± 11%; *P* < 0.001) (**Figure 2F**). Organoids derived from regressing polyps also displayed reduced viability (**Figure 2G**), supporting a model in which organoid morphology reflects the disease state of polyps (**Figure 2H**). This morphological difference was accompanied by increased mRNA expression of cytoskeleton-related (*FLNA*) and tight junction-associated genes (*TJP1*, *WNK4*) in organoids derived from regressing polyps (**Figure 2I**). Expression of these genes was associated with corresponding promoter methylation patterns (**Supplementary Figure 11**).

### Polyp regression is associated with a partially reverted microbiome state

To delineate microbiome variation across tissue states, we analysed normal mucosa (NM; n = 70), progressing (n = 48), and regressing polyps (n = 45) from *APC^1311/+^* pigs. Alpha diversity, quantified as the Shannon effective number of species (exp H′), was comparable across groups (Kruskal–Wallis p = 0.207; **Figure 3A**), indicating preserved community richness and evenness irrespective of lesion trajectory. In contrast, beta diversity based on Bray–Curtis dissimilarity revealed a modest but significant compositional shift (PERMANOVA R² = 2.2%, p = 0.023; **Figure 3B**). Pairwise testing showed that NM differed from both progressing (R² = 1.7%, p = 0.036) and regressing polyps (R² = 1.9%, p = 0.040), whereas the two polyp states were not separable (R² = 1.3%, p = 0.249), suggesting that microbiome restructuring is primarily associated with the transition from normal to neoplastic tissue, independent of subsequent fate.

**Figure 3.**
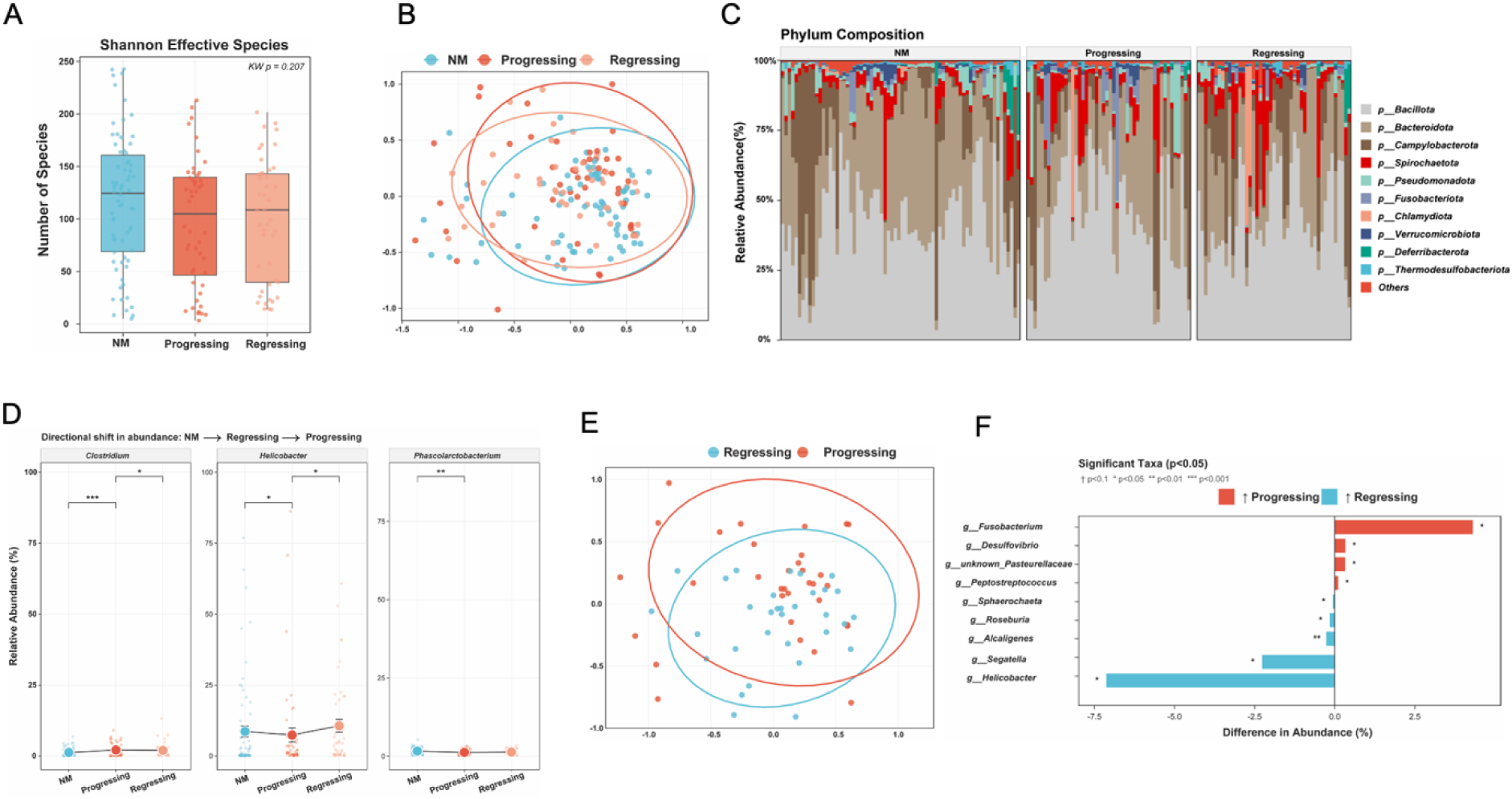
Microbiome composition in progressing and regressing polyps. **(A)** Shannon effective number of species (exp H′) across the three tissue types. Boxes show the interquartile range; horizontal line, median; whiskers, 1.5×IQR; points are jittered individual samples. Kruskal-Wallis p = 0.207. **(B)** Non-metric multidimensional scaling (NMDS) ordination of Bray-Curtis dissimilarity across all three tissue types. Each point represents one sample; ellipses indicate 95% confidence regions. PERMANOVA: R² = 2.2%, p = 0.023. **(C)** Mean phylum-level relative abundance per tissue type. Top 10 phyla are shown; remaining phyla are pooled as “Others” (red). Bacillota and Bacteroidota were the dominant phyla across all groups. **(D)** Mean relative abundance (%) of three key genera across tissue types ordered by polyp status (NM → progressing → regressing). Points show individual samples (jittered); filled circles show group means ± SE; connecting lines illustrate the directional shift across tissue types. Significance brackets indicate pairwise Wilcoxon rank-sum test results (BH-corrected): * p < 0.05, ** p < 0.01, *** p < 0.001. **(E)** NMDS ordination of Bray-Curtis dissimilarity restricted to progressing (n= 33) and regressing (n=32) polyps. PERMANOVA R² = 3%, p = 0.04. **(F)** Differentially abundant genera between progressing and regressing polyps (Wilcoxon rank-sum test, p < 0.05, n = 9 taxa). Bar direction indicates the enriched group. All results are nominal. † p < 0.1, * p < 0.05, ** p < 0.01, *** p < 0.001.

At the phylum level, communities were dominated by Bacillota and Bacteroidota, with Campylobacterota, Spirochaetota, and Fusobacteriota present at low abundance (**Figure 3C**), and no overt compositional shifts detected. Genus-level analysis identified three taxa with nominal differences across tissue types: *Clostridium* (p = 0.004), *Phascolarctobacterium* (p = 0.027), and *Helicobacter* (p = 0.034). *Clostridium* increased stepwise from NM (mean 2.1%) to regressing (2.6%) and progressing polyps (3.7%), whereas *Helicobacter* and *Phascolarctobacterium* showed the opposite trend, declining from NM (16.2% and higher baseline, respectively) through regressing to progressing lesions (**Figure 3D**). Regressing polyps consistently exhibited intermediate abundances, indicative of a partial compositional shift.

Focusing on polyp tissue alone, beta diversity analysis revealed a subtle but significant distinction between progressing and regressing lesions (PERMANOVA R² = 3.0%, p = 0.041; **Figure 3E**). Differential abundance testing identified nine genera with nominal differences (Wilcoxon p < 0.05; **Figure 3F**). Progressing polyps were enriched for *Fusobacterium*, *Desulfovibrio*, *Peptostreptococcus*, and an unclassified Pasteurellaceae genus, whereas regressing polyps showed higher levels of *Helicobacter*, *Segatella*, *Alcaligenes* (p < 0.01), *Roseburia*, and *Sphaerochaeta*. Notably, *Fusobacterium* exhibited the largest effect size, with marked enrichment in progressing lesions, consistent with its established association with colorectal tumorigenesis.

### Single cell atlas of progressing and regressing polyps

To define the molecular, cellular and microbial landscape of polyp fate, we profiled 16 polyps from *APC^1311/+^* pigs by single-cell RNA sequencing (scRNA-seq) (eight matched progressing/regressing pairs; **Figure 4A**). Progressing lesions were classified as high-grade adenomas (n = 8), whereas regressing lesions represented early (n = 4) and late regression (n = 4). Eight samples were processed with 10x Genomics Chromium 3’sequencing and eight with BD Rhapsody, providing complementary capture of epithelial, stromal and immune populations and broad representation of short-lived myeloid cells.

**Figure 4.**
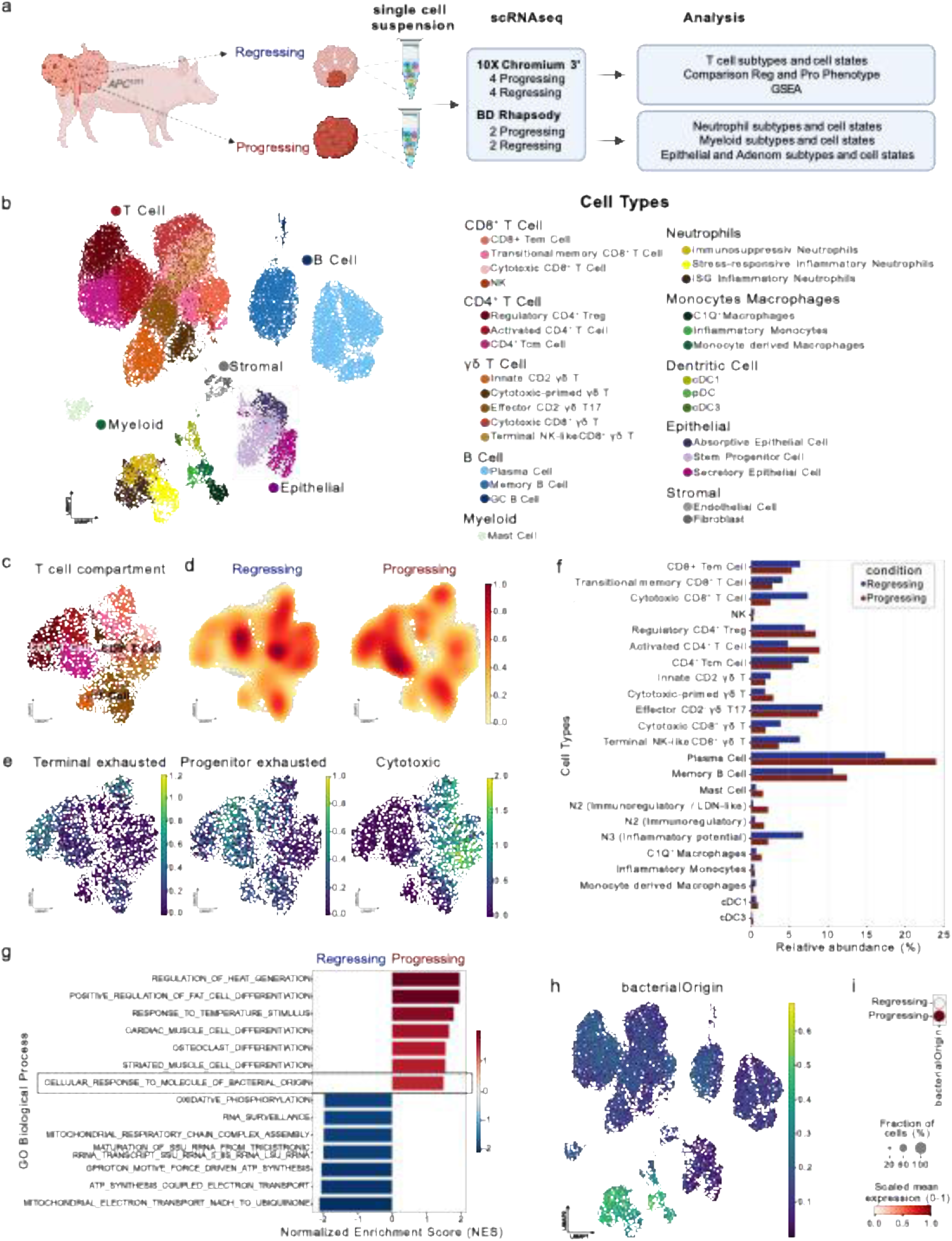
Single-cell RNA-seq resolves immune programs associated with adenoma progression and regression. (a) Experimental setup for scRNA-seq of progressing and regressing polyps from *APC^1311/+^* pigs using 10x Genomics Chromium and BD Rhapsody platforms. (b) UMAP of integrated single-cell profiles showing annotated epithelial, stromal, myeloid, B-cell and T-cell populations. (c) UMAP of the T-cell compartment, including CD4+, CD8+ T-cell subsets. (d) Gaussian kernel density maps showing state shifts between regressing and progressing polyps. (e) Module scores for terminal exhaustion, progenitor exhaustion and cytotoxicity. (f) Relative abundance of immune and stromal/epithelial cell states in progressing and regressing polyps. (g) Gene Set Enrichment Analysis (GSEA) of GO Biological Process terms enriched in progressing or regressing lesions. (h,i) Cellular response to molecule of bacterial origin signatures indicate heightened microbial sensing, particularly in myeloid cells from progressing polyps.

After quality control, UMAP clustering resolved 35 cellular subpopulations across T-cell, B-cell, myeloid, stromal and epithelial compartments (**Figures 4B** and **5**). Because porcine cell-type markers are substantially less comprehensively annotated than human or mouse markers, cluster annotation combined canonical lineage markers, porcine gene annotations, orthologue mapping and iterative marker refinement.

**Figure 5.**
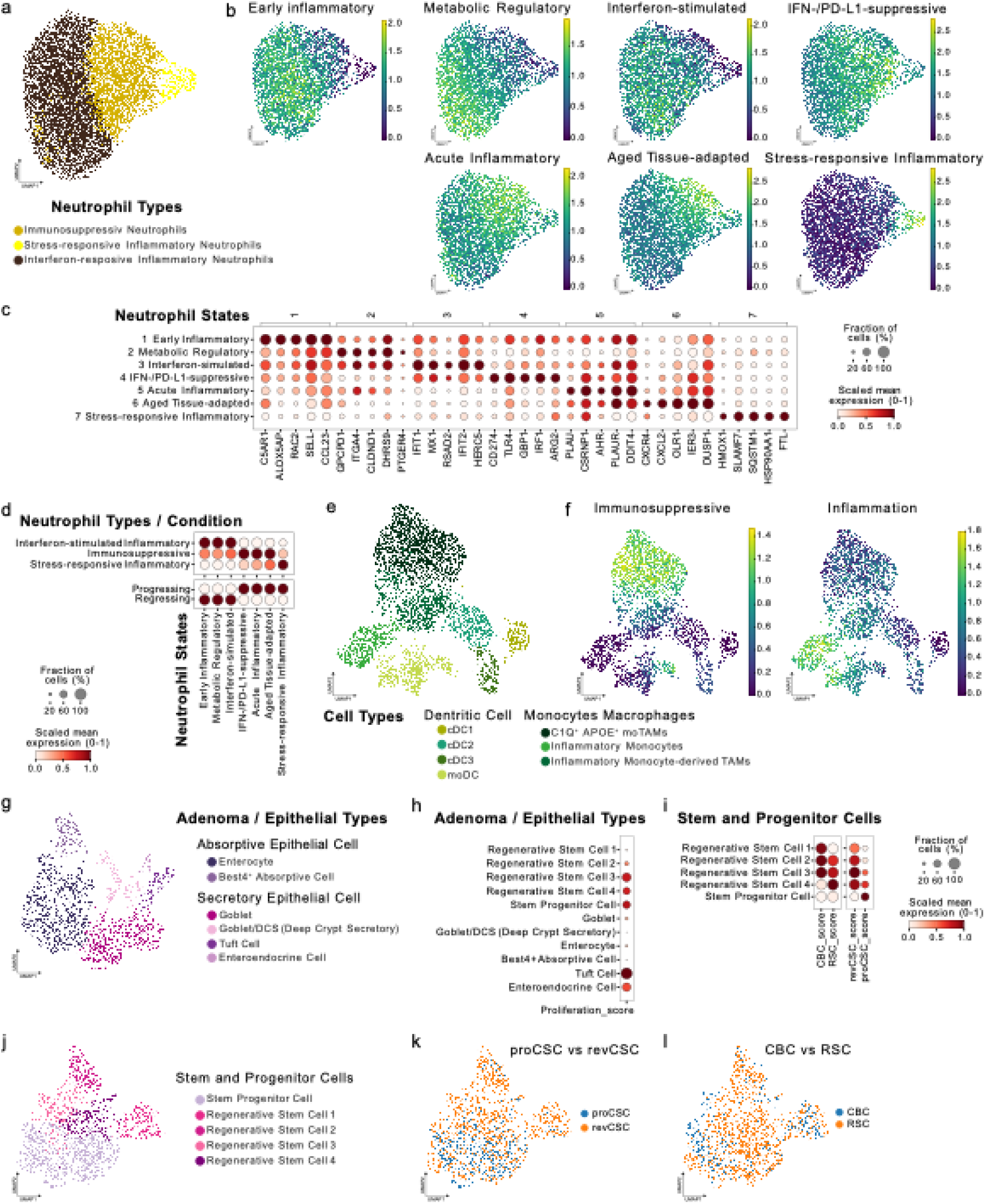
Single-cell analysis identifies distinct myeloid and epithelial states in progressing and regressing polyps. (a) UMAP of the neutrophil compartment showing immunosuppressive, stress-responsive inflammatory and interferon-responsive inflammatory neutrophil populations. (b) Neutrophil module scores for early inflammatory, metabolic regulatory, interferon-stimulated, IFN/PD-L1-suppressive, acute inflammatory, aged tissue-adapted and stress-responsive inflammatory states. (c) Dot plot of differentially expressed genes defining neutrophil states. (d) Mapping of neutrophil states to polyp condition and neutrophil type. (e,f) UMAP and module scoring of dendritic-cell, monocyte and macrophage populations, including immunosuppressive and inflammatory programs. (g) UMAP of epithelial populations, including absorptive and secretory cells. (h) Proliferation scores across epithelial types. (i) Stem/progenitor-state scores, including CBC, RSC, proCSC and revCSC signatures. (j-l) UMAPs and comparative module scores for stem and progenitor compartments.

Lesion fate was associated with a pronounced shift in immune states. Regressing polyps were enriched for central-memory CD4+ T cells, cytotoxic CD8+ T cells and inflammatory neutrophils, consistent with active immune surveillance and local effector function (**Figure 4C-F**). In contrast, progressing polyps showed enrichment of regulatory T cells (Tregs), activated CD4 T cells, plasma cells and immunoregulatory neutrophils, indicating a transition toward immune escape.

Gene-set enrichment analysis supported this interpretation: regressing lesions showed higher cytotoxic and immune-effector programs, whereas progressing lesions were enriched for terminal exhausted T cells, epithelial plasticity, stress-response and cellular responses to molecules of bacterial origin (**Figure 4E-I** and **Figure 5**). The bacterial-response signature was particularly evident in myeloid cells, suggesting that microbial exposure is linked to reprogramming of the local innate immune compartment (**Figure 4H**).

Subclustering of the myeloid compartment resolved immunosuppressive, interferon-responsive inflammatory and stress-responsive inflammatory neutrophil states (**Figure 5A-D**). Progressing lesions contained more immunosuppressive/IFN-PD-L1-associated neutrophils, whereas regressing lesions retained inflammatory neutrophil states. Monocyte, macrophage and dendritic-cell subsets also differed in inflammatory and immunosuppressive scores, supporting coordinated myeloid remodelling during progression (**Figure 5E-F**).

Epithelial subclustering identified absorptive, secretory, adenomatous, stem and progenitor populations (**Figure 5G-L**). Progressing polyps exhibited epithelial plasticity and progenitor-like programs. Together, the single-cell data indicate that regression is associated with a cytotoxic immune-reactive niche, while progression is characterized by epithelial plasticity, myeloid immunosuppression and impaired T-cell effector activity.

### Spatial transcriptomics links bacterial invasion to T cell dysfunktion and neutrophil mediated immunosuppression

Building on the scRNA-seq atlas and 16S microbiome profiling, and the finding that gene expression signatures of bacterial origin are linked with polyp progression, we used Xenium spatial transcriptomics to map host cell states and bacterial signal locatisation within intact polyp architecture (**Figure 6A**). Sixteen formalin-fixed, paraffin-embedded polyps (eight progressing and eight regressing) were profiled with a custom 480-gene host panel and targeted microbial probes, including pan-bacterial 16S EUB plus probes for Epsilonproteobacteria, Fusobacteria, Bacteroides and Firmicutes, enabling host and microbial transcriptomic readouts on the tissue section.

**Figure 6.**
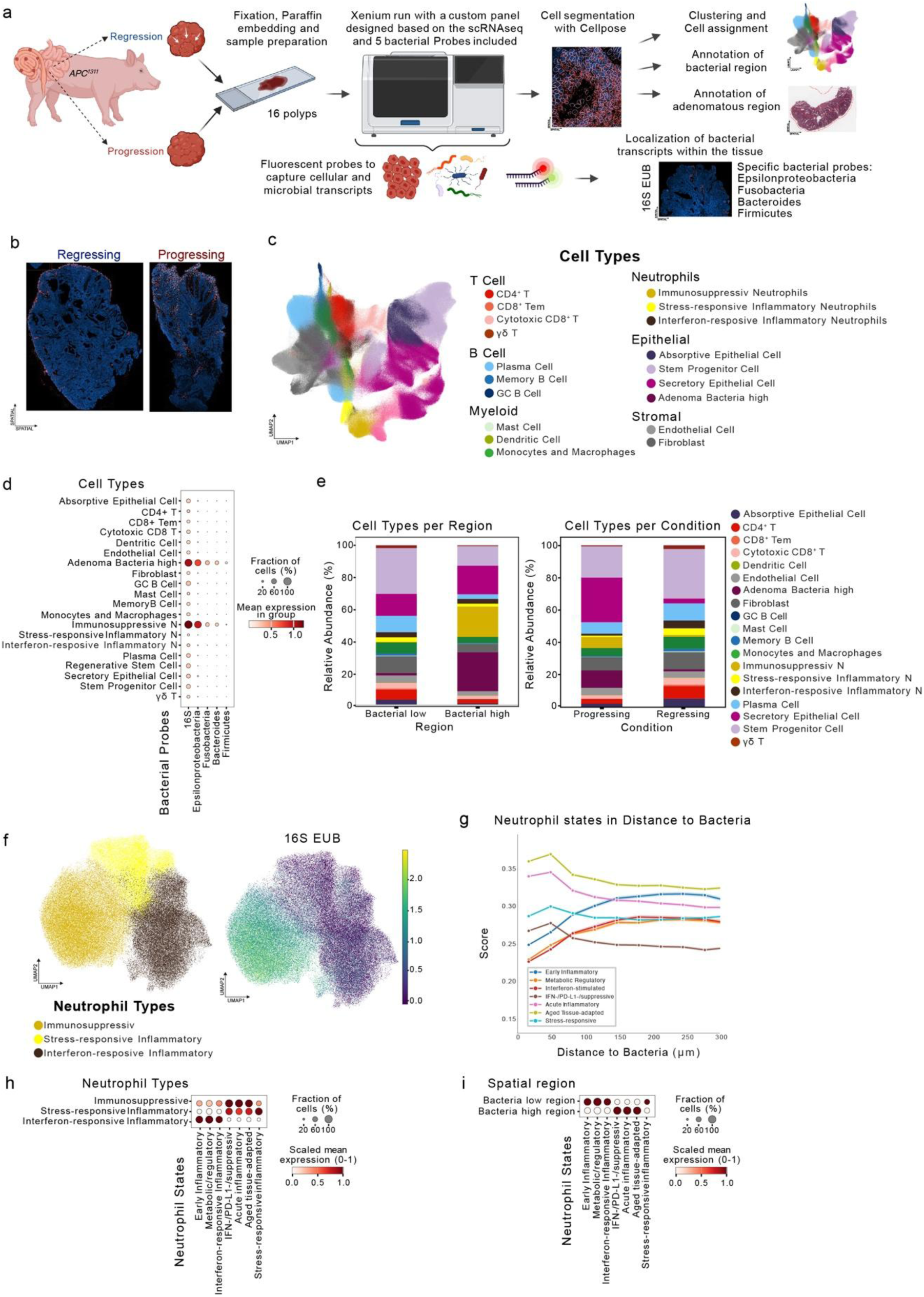
Spatial transcriptomics links bacterial infiltration to cell-type composition and neutrophil state. (a) Overview of Xenium spatial profiling of 16 progressing and regressing polyps from APC1311/+ pigs using a custom host gene panel and targeted bacterial probes for pan-bacterial 16S EUB, Epsilonproteobacteria, Fusobacteria, Bacteroides and Firmicutes. (b) Representative spatial images showing bacterial invasion in progressing lesions and lumen-adjacent bacterial localization in regressing lesions. (c) UMAP of annotated cell types from the spatial experiment. (d) Distribution of bacterial signal across cell types, highlighting enrichment in adenoma cells and immunosuppressive neutrophils. (e) Cell-type composition in bacteria-low versus bacteria-high regions and in progressing versus regressing lesions. (f) UMAP of neutrophil types with bacterial transcript signal. (g) Neutrophil state scores as a function of distance to bacterial signal. (h) Dot plot linking neutrophil states to neutrophil types. (i) Dot plot comparing neutrophil states in bacteria-low and bacteria-high regions.

Spatial analysis resolved microbes (**Figure 6B**) as well as epithelial, stromal, T-cell, B-cell, myeloid and neutrophil populations within tissue context (**Figure 6C**). Comparison of matched H&E and Xenium images showed distinct microbial localization patterns: progressing polyps consistently displayed bacterial invasion into adenomatous regions, whereas regressing polyps showed predominantly lumen-adjacent bacterial signal (**Figure 6B**).

Cell-level quantification revealed that bacterial transcripts were not randomly distributed. Bacterial signal was enriched in adenoma cells and in immunosuppressive neutrophils, and these cell populations formed spatially coupled niches in progressing lesions (**Figure 6D-F**). Bacteria-high regions in progressing polpy showed altered cell-type composition compared with bacteria-low regions, including an increased contribution and abundance of adenomatous cells and immunosuppressive neutrophils and a relative reduction of immune-surveillance-associated cell states including T cells depletion.

Distance-based analyses further demonstrated that neutrophil transcriptional states varied with proximity to bacterial signal. Neutrophils located near bacterial transcripts showed higher immunosuppressive and IFN/PD-L1-associated state scores, whereas bacteria-low or bacteria-distant regions retained more inflammatory/stress-responsive neutrophil programs (**Figure 6G-I**).

These spatial data provide an in situ link between microbial invasion, myeloid reprogramming and T-cell exclusion. Together with the scRNA-seq atlas, they support a model in which bacterial infiltration of adenomatous tissue promotes an immunosuppressive neutrophil niche that facilitates lesion progression, while regression is associated with restricted luminal bacterial localization and spatially organized cytotoxic immune surveillance.

## Discussion

To our knowledge, this study is the first large-scale, longitudinal and multimodal analysis in pigs to link bacterial invasion of early colorectal adenomas with the formation of immunosuppressive, T-cell-excluded niches and adenoma progression. The scale and depth of the dataset are central to this advance. Serial colonoscopy of 82 *APC*^1311/+^ pigs across three generations generated more than 30,000 polyp observations between 3 and 9 months of age. These longitudinal data were integrated with histopathology, organoid assays, regional molecular analyses, mucosal microbiome profiling, an approximately 200,000-cell scRNA-seq atlas of eight matched progressing-regressing polyp pairs, and Xenium profiling of 16 intact polyps using a 480-gene host panel and targeted bacterial probes. Together, this represents an unusually comprehensive large-animal dataset, spanning many independent lesions and millions of cell-level host and bacterial transcript measurements. It permits polyp fate to be resolved at three complementary levels: across animals, across individual lesions and within the microregional cellular niches of each lesion. Previous studies established *APC*^1311/+^ pigs as a genetically and anatomically relevant model of human FAP^12–14^; the present work now uses this model to define how microbial localisation, immune state and epithelial phenotype converge during the earliest stages of colorectal tumour progression.

Longitudinal surveillance revealed that early adenomas do not follow a uniformly progressive trajectory. At baseline, all 136 representative lesions were low-grade adenomas with similar endoscopic morphology, yet between 6 and 9 months many lesions entered a distinct regressive state, whereas others continued to enlarge and, during extended follow-up, reached up to 40 mm. Regressing polyps became smaller, lost crypts and mucin, exhibited lower Ki67 and β-catenin signal, and developed a zonated architecture with epithelial disintegration, crypt abscesses, apoptotic debris and dense immune-cell infiltration. Increased caspase-3, TUNEL and Annexin V positivity further establishes regression as an active process of cell death and tissue clearance rather than passive growth arrest. The coexistence of progressing and regressing lesions in the same genetic background, often within the same animal, therefore exposes a biologically informative decision point at which local environmental and immune factors can override a shared inherited predisposition.

Organoid and regional molecular data show that regression is also accompanied by a distinct epithelial state. Organoids derived from regressing polyps had lower viability, formed more branched and fewer cystic structures, and expressed higher levels of *FLNA*, *TJP1* and *WNK4*, consistent with altered cytoskeletal organisation, epithelial differentiation and junctional control ^24, 25^. Regressing lesions additionally showed a modest reduction in mutant *APC* transcript abundance, increased *APC* copy number and region-specific methylation changes, together with altered proliferation, stem-cell, Wnt, mucin and cytotoxic/apoptotic gene programmes ^26^. These results suggest that epithelial-intrinsic changes contribute to regression and may limit the ability of bacteria to establish a stable intralesional niche. However, epithelial state alone does not account for the strong spatial divergence in immune composition and bacterial localisation. Rather, the data support reciprocal interaction: epithelial plasticity and impaired containment favour bacterial access during progression, whereas epithelial differentiation and immune-mediated tissue clearance accompany bacterial restriction during regression.

A key conceptual finding is that the spatial behaviour of the microbiota was more closely linked to lesion fate than bulk community composition. Alpha diversity did not differ among normal mucosa, progressing polyps and regressing polyps, and beta-diversity effects were significant but small. Even when progressing and regressing polyps were analysed separately, community structure explained only approximately 3% of the variance. Progressing lesions were nevertheless enriched for taxa associated with colorectal tumourigenesis, most notably *Fusobacterium*, as well as *Desulfovibrio* and *Peptostreptococcus*, whereas regressing lesions showed higher abundance of several other genera ^27–29^. These compositional differences identify candidate organisms but also indicate that taxonomy alone is unlikely to determine whether an adenoma grows or regresses. The decisive feature revealed by spatial profiling was whether bacteria remained at the luminal interface or penetrated into the adenomatous compartment.

Single-cell analysis defined the immune states associated with these divergent microbial geographies. Regressing polyps were enriched for central-memory CD4+ T cells, cytotoxic CD8+ T cells and inflammatory neutrophils, together with cytotoxic and immune-effector programmes. In contrast, progressing polyps contained more regulatory and dysfunctional/exhausted T-cell states, plasma cells and immunoregulatory neutrophils, and their myeloid cells showed enhanced responses to molecules of bacterial origin. Neutrophil subclustering further separated inflammatory states from an immunosuppressive, IFN/PD-L1-associated state that was preferentially expanded in progressing lesions. At the same time, progressing adenomatous cells displayed greater plasticity and progenitor-like programmes, whereas cytotoxic and chemoattractant signals were reduced. Thus, progression is not simply associated with increased inflammation; it is associated with a qualitative conversion of inflammation into a myeloid-dominated, T-cell-dysfunctional state. By contrast, regression retains an immune configuration capable of accessing and eliminating lesional cells ^30–35^.

The Xenium data place bacterial invasion at the centre of this conversion. In progressing polyps, bacterial transcripts extended beyond the lumen and were detected within adenomatous regions, where they were enriched in adenoma cells and immunosuppressive neutrophils. Bacteria-high regions contained more adenomatous cells and suppressive neutrophils but fewer T cells and other immune-surveillance-associated states than bacteria-low regions. Moreover, neutrophils located closest to bacterial transcripts had the highest immunosuppressive and IFN/PD-L1-associated scores, whereas bacteria-distant neutrophils retained inflammatory and stress-responsive programmes. These convergent cell-composition and distance analyses support a model in which bacterial invasion creates spatially restricted immunosuppressive, T-cell-excluded niches. Within these niches, microbial sensing reprogrammes neutrophils towards a PD-L1-associated suppressive phenotype, reduces local cytotoxic T-cell access or persistence, and permits plastic adenomatous epithelial cells to survive and expand. In regressing polyps, by contrast, bacterial signal remained predominantly lumen-adjacent and did not form the intralesional bacteria-high niches observed in progressing lesions; inflammatory neutrophils and cytotoxic lymphocytes were maintained, and adenomatous cells underwent apoptosis and structural collapse.

These observations suggest a feed-forward bacteria-neutrophil-epithelium circuit in early tumour progression. Initial epithelial remodelling may permit microbial penetration; bacteria then recruit or reprogramme myeloid cells; immunosuppressive neutrophils dampen cytotoxic T-cell activity; and reduced immune pressure allows further epithelial plasticity, persistence and tissue disorganisation, thereby sustaining bacterial access. The opposite trajectory is observed when bacterial penetration is contained: inflammatory innate responses and cytotoxic lymphocytes remain functionally coupled to the lesion, leading to elimination rather than persistence. This framework extends previous observations that FAP mucosa can harbour tumourigenic bacterial biofilms, that *Fusobacterium nucleatum* accelerates intestinal tumourigenesis and alters the myeloid microenvironment, and that intratumoural bacteria can occupy immunosuppressive microniches associated with myeloid recruitment and T-cell exclusion ^36–38^. The present study advances those findings by demonstrating this architecture in early adenomas rather than established cancers, across multiple independently sampled lesions in a clinically relevant large-animal model, and by directly linking it to longitudinally defined progression and regression states.

Several limitations define the next steps. The spatial and single-cell data establish a strong, internally consistent association but do not yet prove that bacterial invasion is necessary or sufficient for progression, nor whether neutrophil reprogramming is the initiating event or part of a reinforcing loop. The targeted bacterial panel resolves broad taxonomic groups rather than strain-level virulence or metabolic function, and spatial profiling was performed at selected endpoints rather than repeatedly in the same lesion. Direct perturbation will therefore be essential, including defined colonisation, selective microbial depletion, barrier-modifying interventions and manipulation of neutrophil recruitment or PD-L1/CSF1-associated programmes. Validation in human FAP and sporadic adenomas will also be needed, as will assessment under Western-diet and inflammatory conditions. Nonetheless, the data identify bacterial localisation as a potentially more informative biomarker than bulk abundance and nominate the bacterial invasion-neutrophil immunosuppression-T-cell exclusion axis as a tractable point for colorectal cancer interception. Taken together, our findings support a model in which inherited *APC* dysfunction establishes adenoma susceptibility, but bacterial entry into adenomatous tissue reshapes the local immune ecosystem and biases the lesion towards progression; containment of bacterial invasion preserves cytotoxic immune surveillance and permits regression.

## Conclusion

The central conclusion is that bacterial invasion of the adenoma, rather than dysbiosis alone, is linked to progression. In progressing lesions, bacteria penetrate adenomatous regions and create microregional niches enriched for immunosuppressive IFN/PD-L1-associated neutrophils, depleted of T cells and other immune-surveillance states, and coupled to epithelial plasticity. These spatially organised, T-cell-excluded niches provide a mechanism by which microbial access can disable local immune control and permit adenomatous cells to persist and expand. Regressing lesions show the opposite architecture: bacterial signal remains predominantly lumen-adjacent, inflammatory neutrophil and cytotoxic T-cell programmes are preserved, and lesional cells undergo apoptosis and structural clearance.

These findings establish the bacterial invasion-neutrophil immunosuppression-T-cell exclusion axis as a defining feature of early adenoma progression in the porcine FAP model. They further suggest that the depth and location of bacterial penetration may be more informative than bulk microbiome composition for predicting lesion behaviour. Preventing bacterial entry, restoring epithelial containment or disrupting bacteria-conditioned suppressive neutrophil states may therefore offer strategies to redirect early adenomas from progression towards immune-mediated clearance.

Although causal perturbation and human validation remain required, the convergence of longitudinal, molecular, single-cell, microbiome and spatial evidence provides a strong mechanistic framework: *APC* mutation creates a permissive epithelial field, whereas bacterial invasion establishes an immunosuppressive, T-cell-excluded niche that is linked to adenoma progression. This model defines a testable route for early colorectal cancer interception and positions the *APC*^1311/+^ pig as a powerful platform for evaluating microbiome-and immune-directed prevention strategies.

## Writing assistance

None

## Author contributions

TF, KF, DS, AS, FE, DH contributed to the study conception and design. WL, LF, TW, QCh, TMP, DS performed the experiments. WL, LF, DL, PP, FS, YZ, GZ, MM, FE, DH, DS, KF analysed and interpreted data. WL, AS, DS, KF, TF wrote the paper. All authors provided active and valuable feedback on the manuscript.

## Data availability statement

All data relevant to the study are included in the article.

