## Supplementary material for "Microbial Invasion and Immunosuppression Drives Adenoma Progression in Early Colorectal Cancer Development": suplementary table 1

[5] EMBL Heidelberg

[6] Department of Ophthalmology, Columbia University Irving Medical Center, NY 10032, USA

[7] Department of Rehabilitation and Regenerative Medicine, Department of Microbiology and Immunology, Columbia University Irving Medical Center, New York, NY, USA

[8] Leiden University Medical Center, NL

[9] Centre for Biotechnology and Biomedicine, University of Leipzig, Leipzig, Germany

[10] Chair of Livestock Biotechnology, School of Life Sciences, Technische Universität München, 85354 Freising, Germany

[11] Chair of Reproductive Biotechnology, School of Life Sciences, Technische Universität München, 85354 Freising, Germany

### These authors contributed equally to this work.

* These senior authors contributed equally to this work.

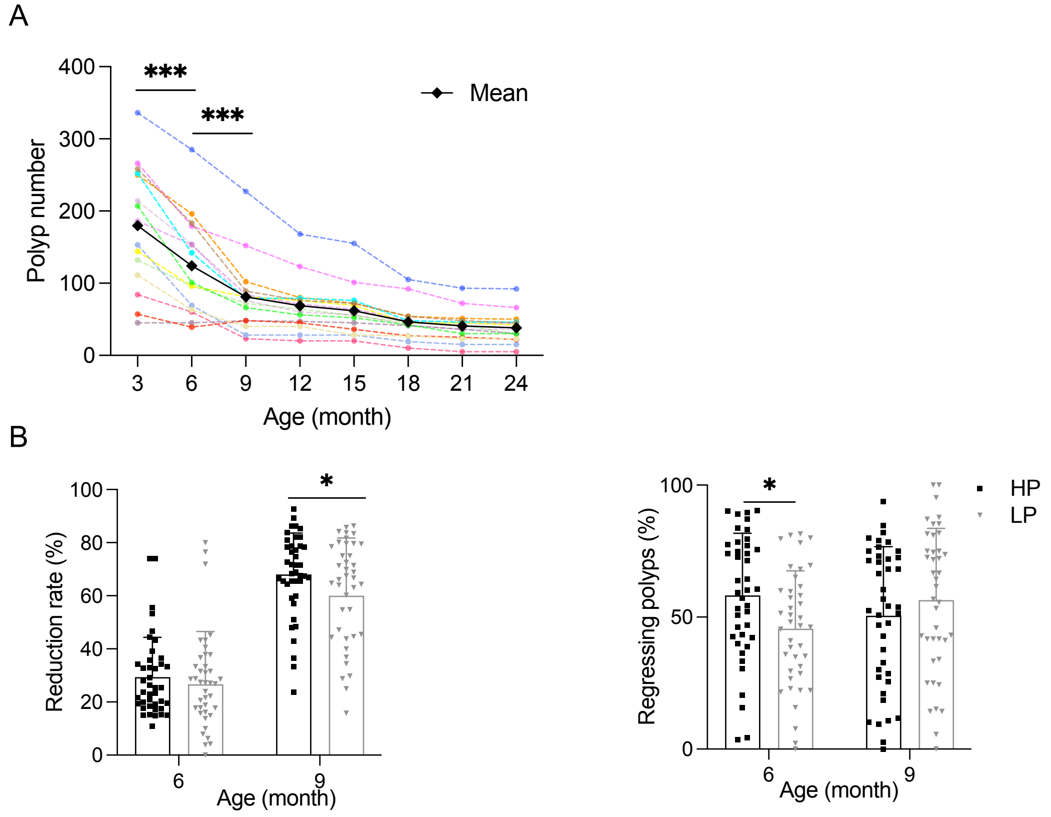


**Supplementary Figure 1.** (A) Number of polyps observed from individual *APC*$^{1311/+}$ pig (n=15) in longitudinal colonoscopy examination for 24 months. (B) Polyp reduction rate (%) and proportion of regressing polyps in HP and LP pigs at the age of 6- and 9- months. The reduction rate was calculated as follows: (polyp number at current colonoscopy - polyp number in previous colonoscopy) / polyp number in previous colonoscopy x 100. Regressing polyps (%) = number of regressing polyps / total number of intestinal polyps x 100.

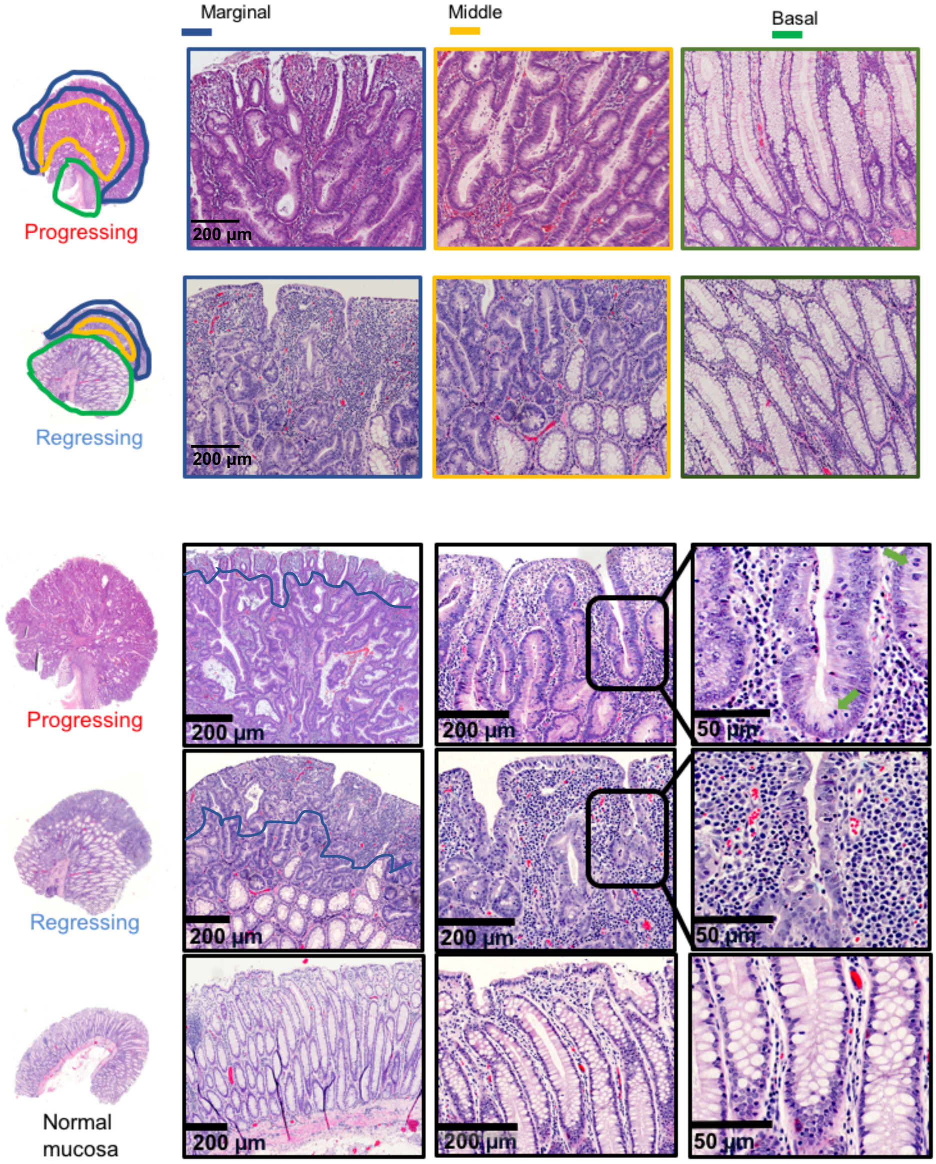
Marginal
Middle
Basal
Progressing
200 µm
Regressing
200 µm
Progressing
200 µm
200 µm
50 µm
Regressing
200 µm
200 µm
50 µm
Normal
mucosa
200 µm
200 µm
50 µm

**Supplementary Figure 2.** A) Three regions which classified basing on histological characteristics in polyps, including blue line circled marginal region, yellow line circled middle region and green line circled basal region. Histological structure of polyps and normal mucosa in different magnification (10x, 40x, 160x). Blue line: the separation boundary between marginal (above the line) and middle region (below the line). Green arrows: meiotic cells.

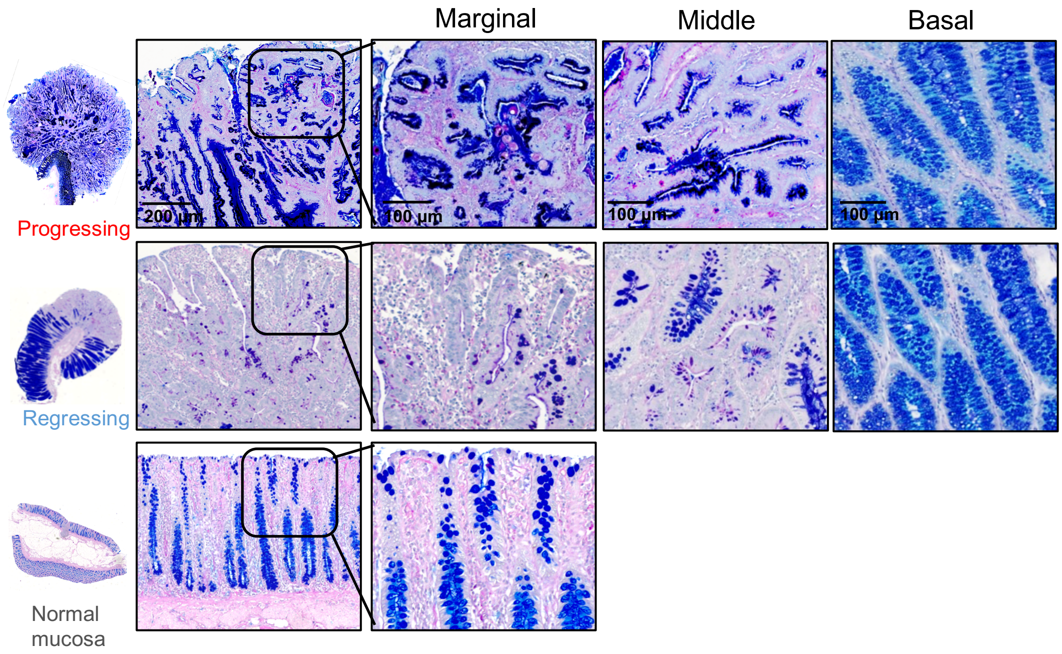


**Supplementary Figure 2.** B) PAS-AB staining of normal mucosa and 3 regions, including marginal, middle and basal region of polyps in *APC*$^{1311/+}$ pigs.

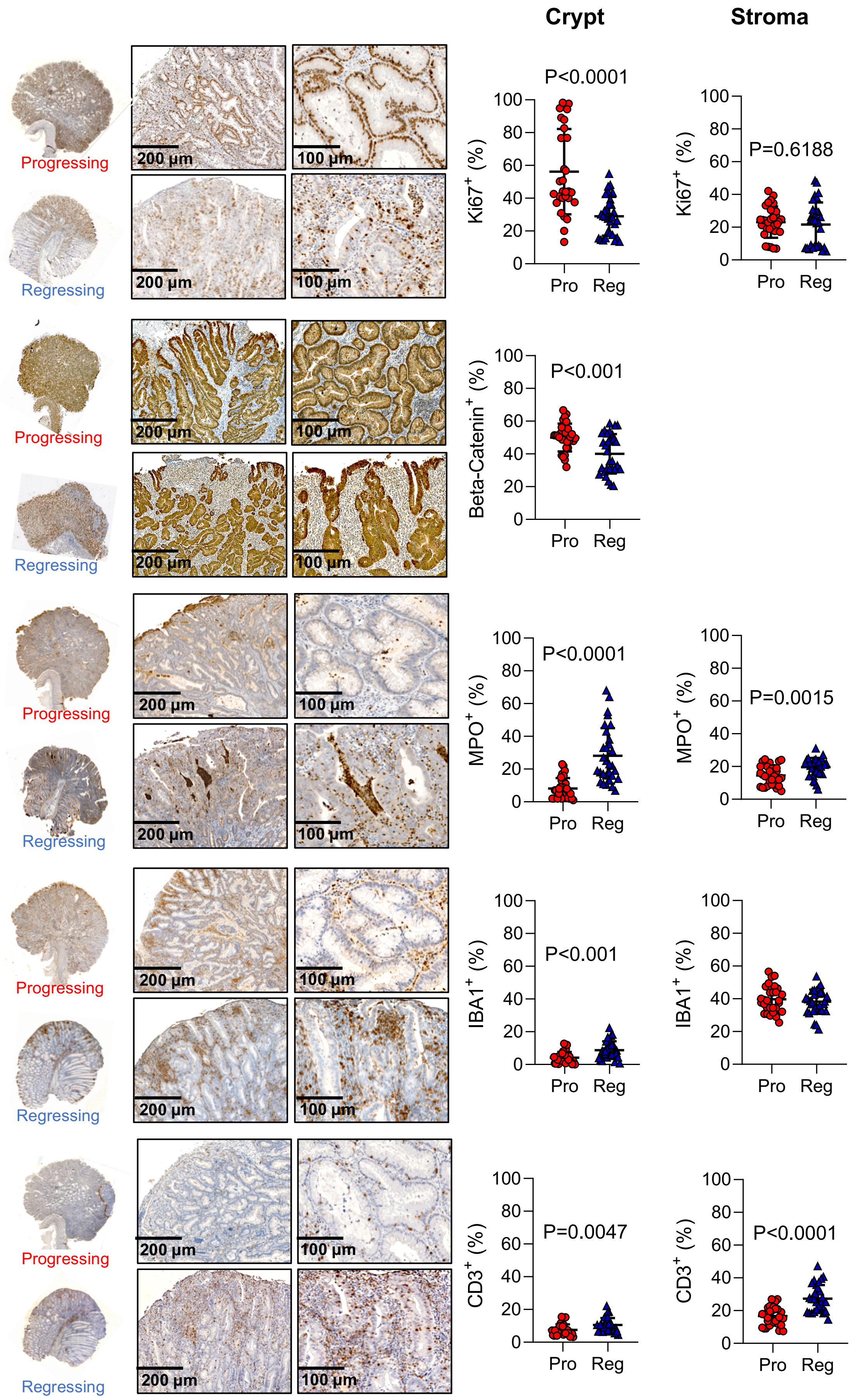
Crypt
Stroma
Progressing
Regressing
200 µm
100 µm
P<0.0001
P=0.6188
Ki67+ (%)
P<0.001
Beta-Catenin+ (%)
P<0.0001
P=0.0015
MPO+ (%)
P<0.001
IBA1+ (%)
P=0.0047
P<0.0001
CD3+ (%)
Pro
Reg

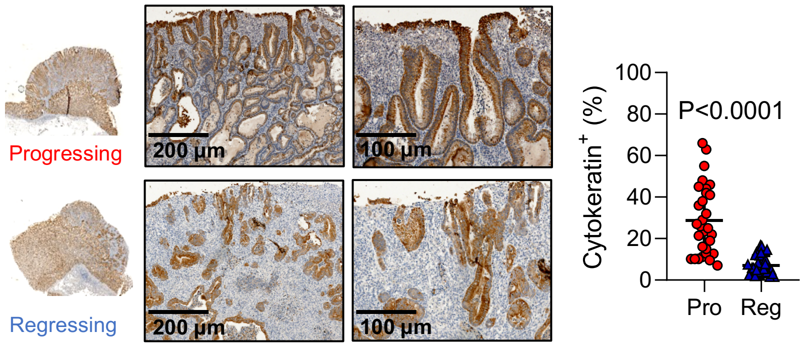


**Supplementary Figure 2**. C) IHC staining for markers of interest and quantitative analysis. IHC for proliferation marker – Ki67, the subunit in intracellular signal transducer of Wnt signaling pathway - Beta-Catenin (D), neutrophil – MPO (E), macrophage - IBA1 (F), T cell - CD3 (G) and cytoskeleton -Cytokeratin (Pan).

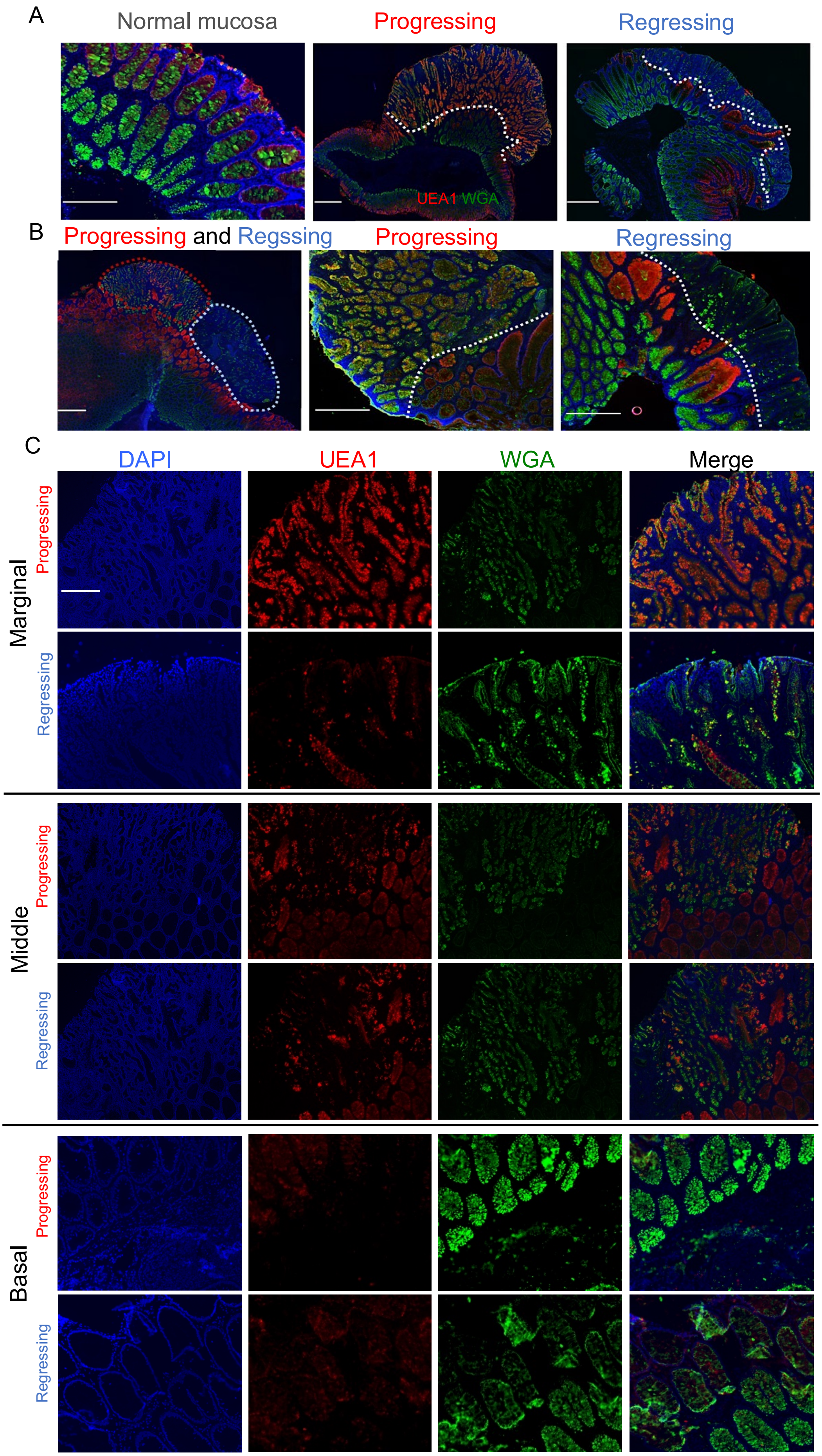
A
Normal mucosa
Progressing
Regressing
UEA1 WGA
B
Progressing and Regssing
Progressing
Regressing
C
DAPI
UEA1
WGA
Merge
Marginal
Progressing
Regressing
Middle
Progressing
Regressing
Basal
Progressing
Regressing

**Supplementary Figure 3**. Representative pictures of UEA1 and WGA staining for normal mucosa and polyps in *APC*$^{1311/+}$ pigs. (A) Representative pictures for normal mucosa (left) and progressing (middle) and regressing polyps (right). (B) Representative pictures of UEA1 and WGA staining for progressing and regressing polyps. Left: picture of a progressing polyp (red circle) next to a regressing polyp (light blue circle) in the same section. Middle: different UEA1+ staining in adenomatous lesion and normal mucosa-like structure of progressing polyp as separating line (white dotted line) with in 10x microscopic power field. Right: different UEA1+ staining in adenomatous lesion and normal mucosa-like structure of regressing polyp as separating line (white dotted line) with in 10x microscopic power field. Scale bar: 200 μm, Red stain: UEA1+ signal, green stain: WGA+ signal, blue stain: DAPI. Scale bar: 200 μm. **(C)** Pictures from single channel of DAPI, UEA1 (TRIRC) and WGA (FITC) staining from marginal, middle and basal region of progressing and regressing polyps in 10x microscopic power filed. Scale bar: 200 μm.

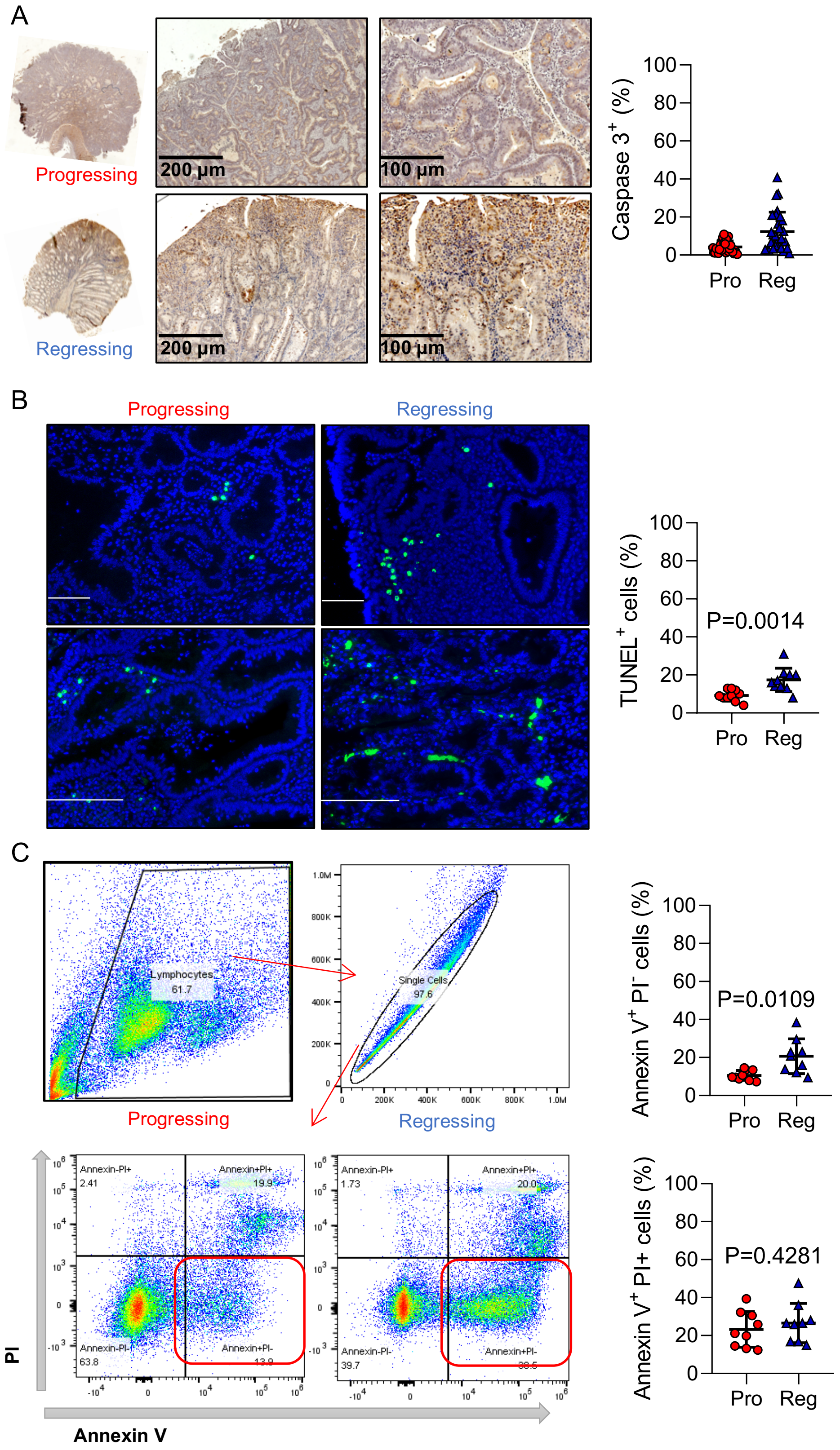


**Supplementary Figure 4**. (A) IHC for apoptotic marker – caspase 3. (B) Staining (left) and statistics (right) of apoptotic cells by TUNEL assay in progressing and regressing polyps. Cell nucleus stained in blue (DAPI) and apoptotic cells stained in green (FITC). Bar=100 μm. (C) Flow cytometry result (left) and analysis (right) of Annexin V and PI staining for apoptotic cells and dead cells in progressing and regressing polyps. Red box: Annexin V+ and PI- cells, apoptotic cells. Grey box: Annexin V+ and PI+ cells, necrotic or dead cells.

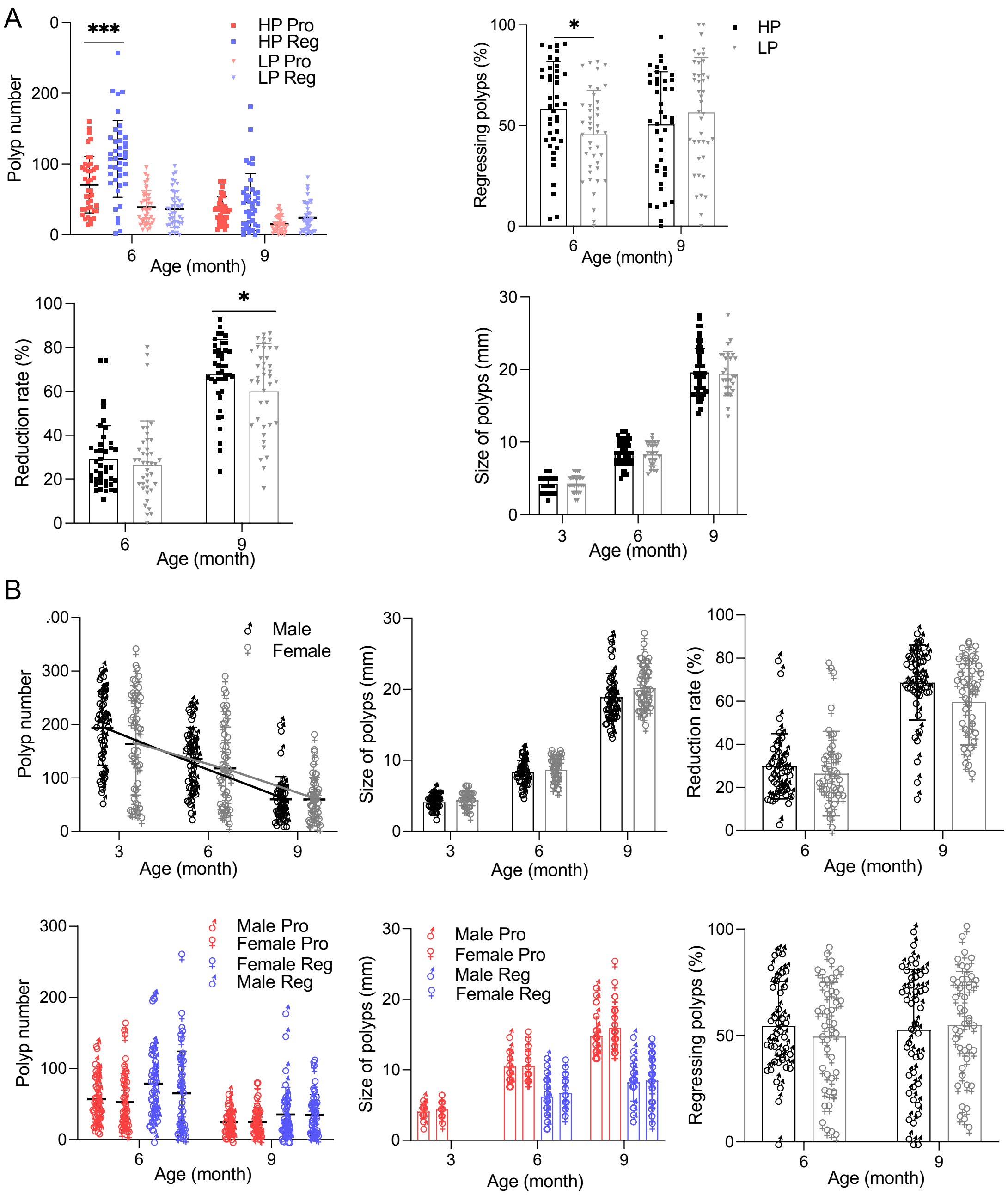
A
***
HP Pro
HP Reg
LP Pro
LP Reg
Polyp number
200
100
0
6
9
Age (month)
*
HP
LP
Regressing polyps (%)
100
50
0
6
9
Age (month)
*
Reduction rate (%)
100
80
60
40
20
0
6
9
Age (month)
Size of polyps (mm)
30
20
10
0
3
6
9
Age (month)
B
Male
Female
Polyp number
300
200
100
0
3
6
9
Age (month)
Size of polyps (mm)
30
20
10
0
3
6
9
Age (month)
Reduction rate (%)
100
80
60
40
20
0
6
9
Age (month)
Male Pro
Female Pro
Female Reg
Male Reg
Polyp number
300
200
100
0
6
9
Age (month)
Male Pro
Female Pro
Male Reg
Female Reg
Size of polyps (mm)
30
20
10
0
3
6
9
Age (month)
Regressing polyps (%)
100
50
0
6
9
Age (month)

**Supplementary Figure 5.** (A) Progressing and regressing polyp numbers in HP and LP group pigs at age of 6 and 9 months (top left). Percentage of regressing polyps in HP and LP group pigs (top right). Reduction rate in HP and LP group pigs at age of 6 and 9 months (bottom left). Average size of polyps in diameter (mm) from HP and LP group pigs at the age of 3, 6 and 9 months (bottom right). (B) Colorectal polyp numbers counted from colonoscopy examination for male and female APC1311/+ pigs at the age of 3 month (3M), 6 month (6M) and 9 month (9M), respectively (top left). Average size of polyps in diameter (mm) from Male and Female group pigs at the age of 3, 6 and 9 months (top middle). Polyp reduction rate in Male and Female group pigs at the age of 6 month and 9 months (top right). The number of progressing and regressing polyps in Male and Female group pigs at the age of 6 and 9 months (bottom left). Average size of progressing and regressing polyps in diameter (mm) from Male and Female group pigs at the age of 3, 6 and 9 months (bottom middle). Proportion of regressing polyps in Male and Female group pigs at the age of 6 and 9 months, Regressing polyps (%) = number of regressing polyps / total number of intestinal polyps * 100 (bottom right).

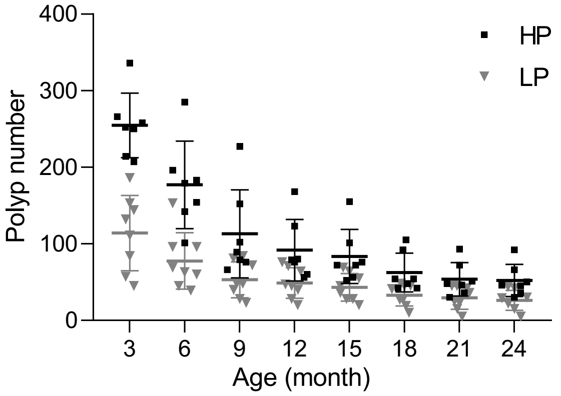


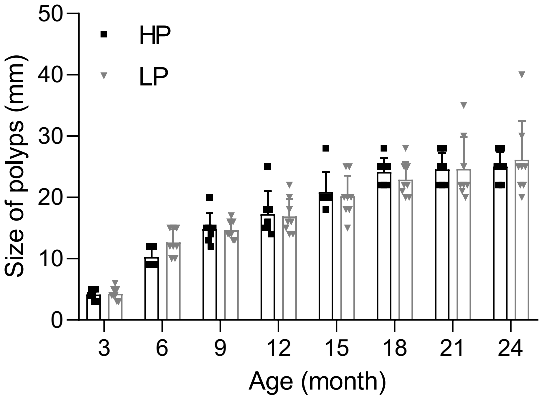


**Supplementary Figure 6.** Colorectal polyp numbers counted from trimonthly longitudinal colonoscopy examination for HP and LP group APC1311/+ pigs from the age of 3 months until 24 months (left). Average polyp size analyzed from trimonthly longitudinal colonoscopy examination for HP and LP group APC1311/+ pigs from the age of 3 months until 24 months (right).

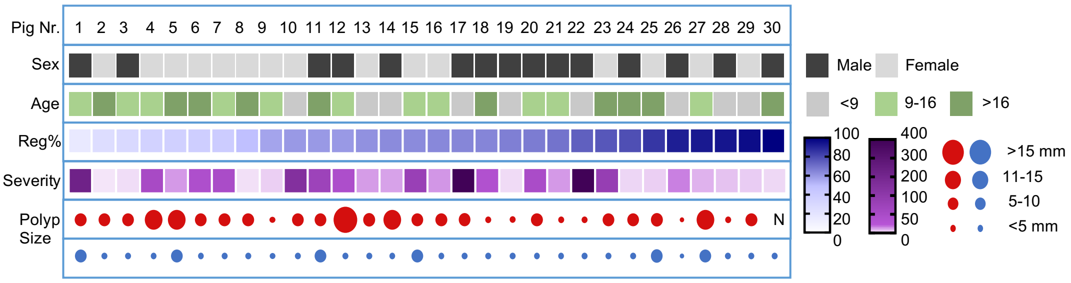


**Supplementary Figure 7.** Basic information of 30 APC1311/+ pigs. Reg% represents the percentage of regressing polyps out of total polyps. Severity is represented by the number of polyps, which ranges from 0-1000. The point size represents the polyp size, red point: progressing polyps, blue point: regressing polyps.

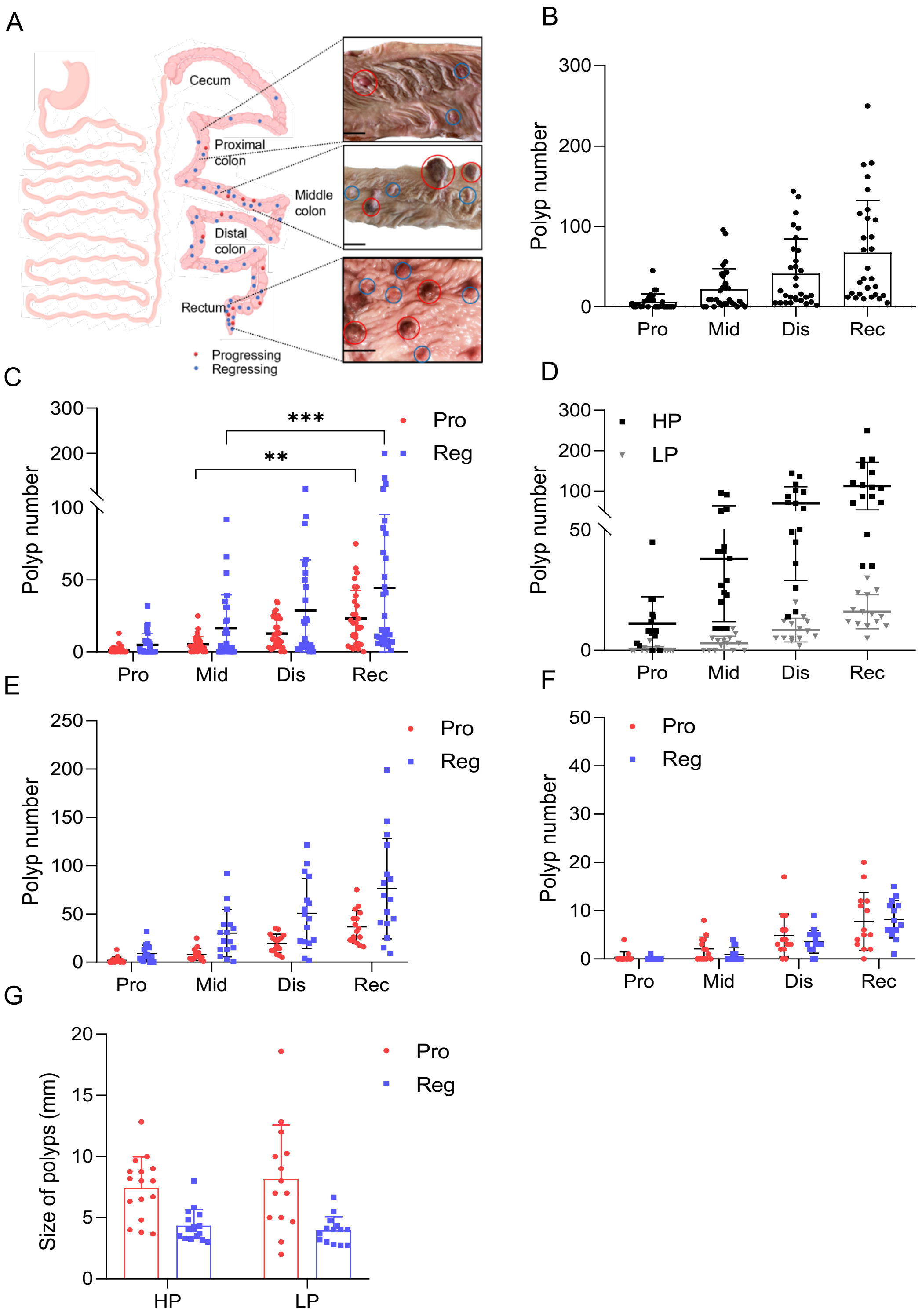


**Supplementary Figure 8.** (A) The examined digestive tract of pig including stomach, small intestine, colon and rectum with progressing and regressing polyps are also schematically shown. The pictures taken from rectum, middle colon and proximal colon with polyps are shown in the right zoom-in panel (red circle: progressing polyp, blue circle: regressing polyp, bar: 1cm). (B) Polyp numbers counted from proximal colon, middle colon, distal colon and rectum. (C) Progressing and regressing polyp number counted from proximal colon, middle colon, distal colon and rectum. (D) Polyp numbers counted from proximal colon, middle colon, distal colon and rectum in HP and LP pigs. (E) Progressing and regressing polyp number counted from proximal colon, middle colon, distal colon and rectum in HP pigs. (F) Progressing and regressing polyp number counted from proximal colon, middle colon, distal colon and rectum in LP pigs. (G) Average size of progressing and regressing polyps in HP and LP pigs. Pro: proximal colon, Mid: middle colon, Dis: distal colon, Rec: rectum.

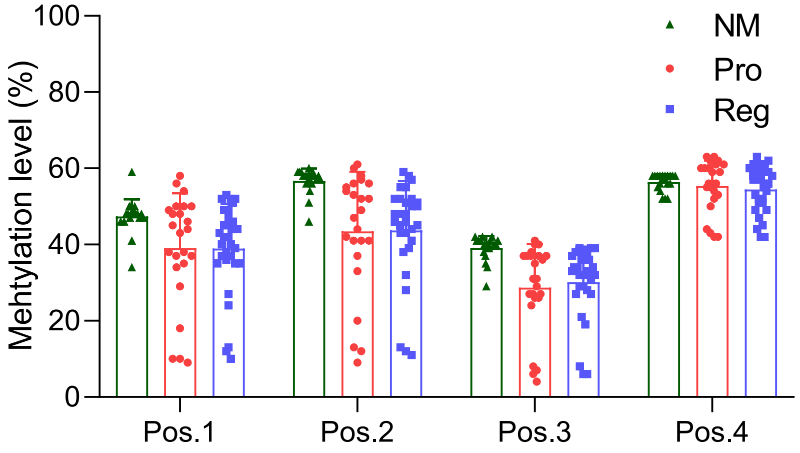


**Supplementary Figure 9.** Methylation level of 4 CpG island position in *APC* gene 3'UTR region.

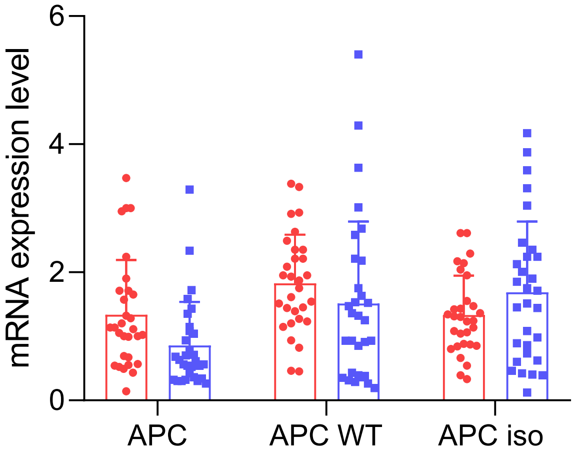


**Supplementary Figure 10.** The mRNA expression of APC with all allele, APC wildtype allele, and APC isoform from bulk polyp samples.

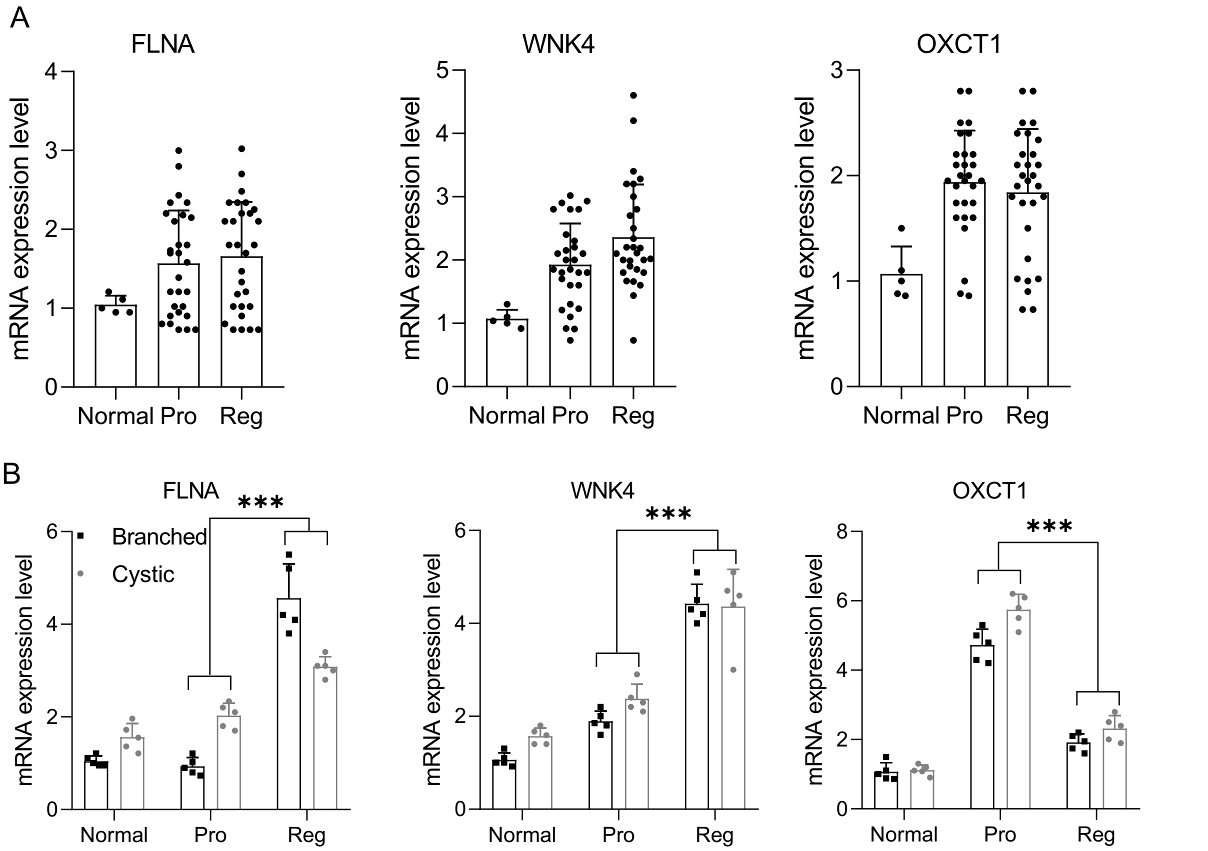


**Supplementary Figure 11.** (A) The mRNA expression level of *FLNA*, *WNK4* and *OCT1* from normal mucosa, progressing and regressing polyps in APC1311 pigs. (B) The mRNA expression level of *FLNA* and *WNK4* in branched and cystic organoids from normal mucosa, progressing and regressing polyps in APC1311 pigs.
